# Endothelial PTBP1 Deletion in Transplanted Cardiac Tissue Limits Cardiac Allograft Vasculopathy

**DOI:** 10.64898/2026.02.18.706637

**Authors:** Christopher L. Pathoulas, Koki Hayashi, Ivy Rosales, Amy L. Kimble, Krish Dewan, Ryan T. Gross, Jenna Lancey, Lifang Ye, Qian Li, Yunfeng Li, Bing Hao, Bo Reese, Evan Jellison, Antoine Menoret, Anthony T. Vella, Dawn E. Bowles, Nicole M. Valenzuela, Jeffrey J. Hsu, Alessandro Alessandrini, Patrick A. Murphy

## Abstract

**Background:** Cardiac allograft vasculopathy (CAV) is a leading cause of late graft failure and mortality following heart transplantation, with limited therapeutic options. Endothelial cells (ECs), at the interface between the donor graft and host immune system, play a central role in CAV development. However, the molecular mechanisms driving endothelial dysfunction and vascular remodeling in chronic heart transplant rejection remain poorly understood.

**Methods:** To characterize endothelial alterations associated with CAV, we isolated nuclei from cardiac tissues of four human donor groups: (1) early post-transplant CAV-negative surveillance biopsies, (2) CAV-negative explanted grafts with acute cellular rejection (ACR), (3) late-stage CAV-positive explanted grafts, and (4) naïve non-transplanted control hearts. We applied intranuclear cellular indexing of transcriptomes and epitopes (inCITE-seq) to profile endothelial gene expression together with nuclear protein levels of splice factor polypyrimidine tract-binding protein 1 (PTBP1), a key post-transcriptional regulator of endothelial inflammatory responses. Functional relevance of PTBP1 was assessed using endothelial-specific deletion of *Ptbp1* in an F1 hybrid murine model of CAV.

**Results:** In human CAV, endothelial cells exhibited increased transforming growth factor-β (TGF-β) signaling and reduced oxidative phosphorylation (OxPhos) transcripts. Nuclear PTBP1 protein levels were markedly elevated in CAV endothelium and were associated with TGF-β-responsive transcriptional programs and correlated with clinical indices of cardiac dysfunction. In murine heart transplants, endothelial-specific deletion of *Ptbp1* markedly reduced hallmarks of CAV, including neointimal hyperplasia, fibrosis, and lymphocyte activation. At the molecular level, endothelial *Ptbp1* deletion prevented suppression of mitochondrial transcripts and preserved mitochondrial content and integrity under hypoxic stress, attenuating interferon signaling in endothelial cells.

**Conclusion:** These findings identify PTBP1 as a central endothelial regulator linking pro-fibrotic stress to mitochondrial dysfunction and immune activation in chronic cardiac allograft rejection. Targeting endothelial PTBP1 may represent a strategy to limit chronic graft injury while minimizing systemic immunosuppression.

## Introduction

Cardiac allograft vasculopathy (CAV), characterized by progressive intimal proliferation within epicardial and intramyocardial vessels, remains a leading cause of late graft failure and mortality in heart transplant (HTx) recipients^1^. Despite advances in immunosuppression that have reduced acute cellular rejection, effective therapies to prevent or reverse CAV development are limited. Moreover, prolonged systemic immunosuppression increases the risk of severe complications, including infection and malignancy. Thus, there is a critical need to identify graft-intrinsic disease mechanisms that can be therapeutically targeted while minimizing systemic immunosuppression.

Endothelial cells (ECs) mediate interactions between donor cardiac tissue and the host immune system and play a central role in CAV development. Graft ECs express HLA, adhesion, and co-stimulatory molecules that promote immune cell activation and recruitment, leading to vascular inflammation and neointimal formation^1^. Pro-inflammatory cytokines, including TNF-α, IFN-γ, and TGF-β, further amplify these responses by priming ECs and inducing a dysfunctional, pro-inflammatory endothelial state^1^. This endothelial dysfunction is central to CAV development and is also a well-established driver of atherosclerosis, which shares many pathological features with CAV, including intimal proliferation and vascular remodeling^2^.

In both conditions, TGF-β signaling plays a prominent role, and direct or indirect perturbation of endothelial TGF-β signaling limits intimal proliferation, immune activation, and cardiac fibrosis in murine models of atherosclerosis and CAV^3–5^. TGF-β contributes to vascular fibrosis through multiple mechanisms including endothelial activation, endothelial-to-mesenchymal transition (EndoMT), and metabolic rewiring^4,6,7^. Recent studies have further linked TGF-β signaling to endothelial mitochondrial dysfunction, in part through promotion of mitochondrial fission, a process associated with increased endothelial immunogenicity and atherosclerosis development^8–10^. Consistent with this, inhibition of endothelial mitochondrial fission attenuates atherosclerosis, heart transplant rejection, and neointima formation^8–10^. However, the cellular mechanisms through which TGF-β and other inflammatory cues reprogram endothelial mitochondrial metabolism to promote vascular remodeling in chronic heart transplant rejection remain poorly defined.

Post-transcriptional gene regulation by RNA-binding proteins (RBPs) plays a key role in controlling cellular metabolism and stress responses. In ECs, the splice factor polypyrimidine tract-binding protein 1 (PTBP1) is an important regulator of alternative splicing of inflammatory and metabolic genes in response to cellular stress and pro-inflammatory stimuli^11,12^, and plays a role in cardiac development^13^. We have previously shown that PTBP1 is induced in the endothelium during atherosclerosis development in a platelet-dependent manner and is required for endothelial “priming,” enhancing responsiveness to TNF-α stimulation *in vitro*^11^. Endothelial PTBP1 is also upregulated in pulmonary arterial hypertension (PAH) where it promotes alternative splicing of *Pkm* towards the PKM2 isoform, shifting metabolism toward glycolysis and away from oxidative phosphorylation^14^. Importantly, endothelial-specific PTBP1 knockout limited atherosclerotic intimal formation and plaque inflammation *in vivo* ^11^. Together, these findings suggest that PTBP1 may act as a molecular switch, reinforcing pro-inflammatory and metabolic endothelial reprogramming.

Here, we applied inCITE-seq multi-omics profiling to human heart transplant samples, comparing late-stage CAV-positive explants with early-stage CAV-negative biopsies. We demonstrate that PTBP1 is elevated in CAV endothelium, where it correlates with enhanced TGF-β signaling, decreased oxidative phosphorylation-related transcripts, and cardiac dysfunction. Using a murine model of CAV, we further show that endothelial deletion of *Ptbp1* prior to transplantation attenuates CAV-associated alterations in oxidative phosphorylation-related gene expression and markedly limits intimal hyperplasia, fibrosis, hypoxia, and local immune activation compared to wild-type grafts. Endothelial *Ptbp1* deletion also preserves mitochondrial content and integrity in human and murine ECs under hypoxic stress, attenuating interferon signaling. Collectively, these findings identify post-transcriptional control of endothelial metabolism as a mechanism linking pro-fibrotic TGF-β signaling to chronic cardiac allograft rejection.

## Results

### Single-nucleus analysis of human heart transplant tissues reveals progressive endothelial cell dysfunction

Heart transplant recipients undergo serial endomyocardial biopsies as part of routine rejection surveillance, providing longitudinal insight into early-stage post-transplant changes (1-3 years). CAV-positive cardiac tissues are typically obtained at the time of re-transplantation (∼10-15 years post-transplant), representing late-stage disease. To define endothelial alterations associated with CAV, we performed intranuclear cellular indexing of transcriptomes and epitopes (inCITE-seq), a multi-omics approach that enables simultaneous quantification of nuclear protein levels using oligonucleotide-tagged antibodies together with single-nucleus RNA sequencing^15^.

Using this approach, we analyzed nuclei isolated from four groups of cardiac tissues from nineteen patients: (1) nineteen early post-transplant CAV-negative surveillance biopsies from five donors, (2) CAV-negative explanted grafts from two donors with moderate acute cellular rejection (ACR) (ISHLT grade 2R), (3) late-stage CAV-positive explants from ten donors, and (4) naïve (non-transplanted) heart tissue from two donors with non-ischemic cardiomyopathy (Figure 1A, SI Table 1).

**Figure 1.**
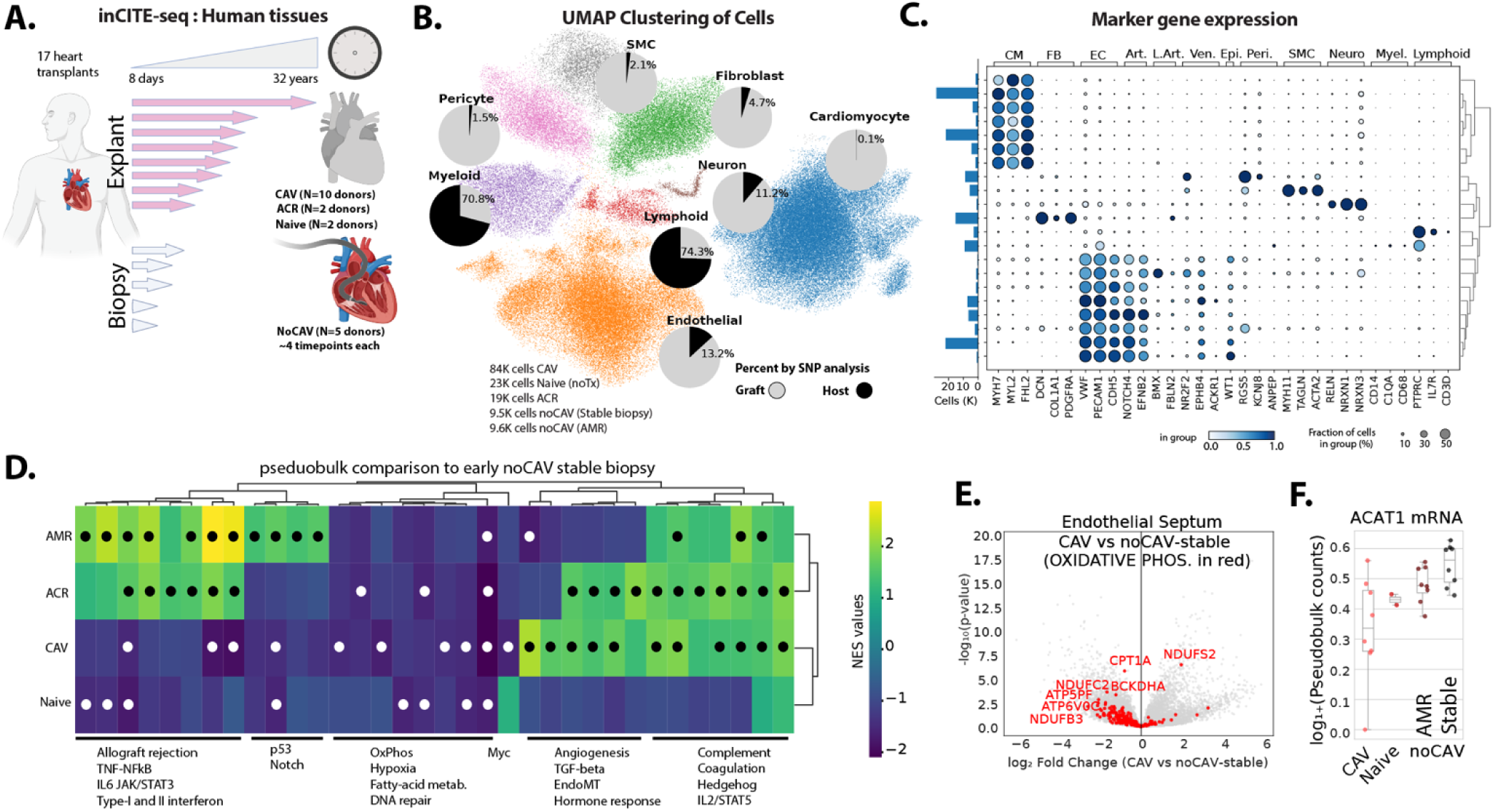
Progressive transcriptional alterations in the endothelium in progression from stable biopsy to allograft vasculopathy. (A) Outline of approach to examine endothelial alterations in cardiac allograft vasculopathy, indicating the source of tissues, routine endomyocardial biopsy or endpoint heart explant. CAV-negative biopsy samples are septal, but both septal and ventricle samples were obtained from CAV-positive explants. (B) Uniform Manifold Approximation and Projection (UMAP) clustering of cells from all combined samples, showing major general cell populations. Inset pie graphs show the percent of each cell type contributed by graft or host, by de novo and reference-free Souporcell analysis of SNP variants. (C) Dot plot showing expression levels of marker genes for general cell populations. (D) Heat map showing enriched MSigDB Hallmark pathways (ranked gene expression, by DESeq2 stat, from pseudobulk analysis), with all comparisons to stable biopsy samples without development of CAV. Main components of each cluster are shown below. Circles = padj < 0.05 (E) Volcano plot showing altered gene expression in the endothelial cell clusters (pseudobulk) in septal samples from CAV-positive explant versus CAV-negative biopsy, with overlay of genes associated with the specific OxPhos MSigDB Hallmark pathway. (F) Targeted analysis of example transcripts showing differential expression, in quartile normalized read counts in each endothelial sample. CAV-negative biopsies were further separated into samples with antibody mediated rejection (AMR) or without signs of antibody or cellular rejection (Stable).

CAV-negative biopsies and CAV-positive explant tissues were obtained from the ventricular septum for direct comparison. In addition, tissues from the ventricular wall were analyzed from a subset of the CAV-positive explants. Early-stage CAV-negative biopsies were further stratified by clinical course; patients #1-3, who had no history of moderate rejection, were classified as stable, whereas patients #4-5 developed antibody-mediated rejection (AMR) during follow-up.

Across all samples, we profiled approximately 126,000 nuclei from explant tissues (84,000 CAV, 19,000 acute rejection without CAV, and 23,000 naïve) and 19,000 nuclei from CAV-negative biopsies (9,500 stable and 9,500 with AMR, Figure 1A-C). Endothelial nuclei comprised ∼25% of nuclei in septal tissues and ∼50% of nuclei following endothelial enrichment from ventricular tissues (Figure 1A&B and SI Figure 1). Prior studies suggest that host-derived endothelial cells can repopulate cardiac grafts^16,17^. To distinguish graft- and host-derived nuclei, we used SNP-based genetic profiling (Figure 1B and SI Figure 2). The endothelium was predominantly graft-derived, with two exceptions showing substantial host-derived endothelial contribution (SI Figure 2).

To define endothelial transcriptional responses associated with transplantation, we compared endothelial cells from biopsy and explant tissues with naïve non-transplant control hearts. Relative to naïve hearts, all the transplanted tissues exhibited shared transplant-associated gene signatures, including enrichment of hypoxia, complement, coagulation, and TNF-α signaling pathways (SI Figure 3 and SI Table 3), indicating persistent inflammatory and stress signaling across graft stages.

Next, we distinguished transcriptional changes associated with acute alloimmune injury from those linked to chronic CAV. Endothelial cells from CAV-negative biopsy samples experiencing AMR showed increased interferon and TNF-α signaling relative to stable biopsies (Figure 1D), consistent with acute inflammatory activation. In contrast, endothelial cells from late-stage CAV-positive explants exhibited a distinct transcriptional profile characterized by increased TGF-β signaling, endothelial-to-mesenchymal transition (EndoMT), and significant suppression of oxidative phosphorylation (OxPhos)-associated transcripts, including *ACAT1*, relative to CAV-negative stable biopsies (Figure 1D-F). Interestingly, endothelial cells from CAV-negative ACR explants exhibited features of both AMR and CAV, including inflammatory activation alongside pro-fibrotic and metabolic signatures (Figure 1D). These findings are consistent with the possibility that moderate ACR induces vascular changes that may represent a transitional state within the microvasculature between acute injury and chronic vasculopathy.

We performed similar analyses across additional cardiac cell populations (SI Table 2, SI Figure 4&5, SI Table 3A&B), revealing notable cell type-specific responses. Whereas OxPhos-related pathways were suppressed in CAV endothelial cells relative to CAV-negative stable tissues, these same pathways were increased in myeloid cells, highlighting diverging metabolic rewiring across cellular compartments during CAV development.

Together, these data indicate that CAV is associated with progressive endothelial dysfunction marked by pro-fibrotic signaling and impaired transcription of genes associated with mitochondrial OxPhos.

### Endothelial PTBP1 is elevated across the CAV vasculature and correlates with cardiac dysfunction

Elevated endothelial PTBP1 expression is associated with chronic vascular remodeling^11^; however whether PTBP1 is increased in the CAV microvasculature has not been defined. To quantify nuclear PTBP1 protein levels and correlate protein expression patterns with endothelial transcriptional states across disease stages, we assessed nuclear PTBP1 by inCITE-seq (Figure 2A). Histone H3 and non-specific IgG were included as positive and negative controls, respectively. Histone H3 was detected in nearly all nuclei (SI Figure 6) and was used to normalize PTBP1 expression, as we have previously reported^15^. Endothelial PTBP1 levels were widely distributed within each sample and elevated in CAV-positive explants compared with CAV-negative biopsies and naïve controls (Figure 2B&C). These changes in endothelial PTBP1 expression were independently validated by immunofluorescence in a subset of the same samples. PTBP1 staining within the lectin-stained vasculature showed a heterogeneous distribution and was significantly increased in CAV-positive samples compared with naïve control tissues (Figure 2D&E), corroborating our inCITE-seq findings. Notably, among CAV-positive explants, the sample derived from a patient whose graft function was maintained for more than 32 years prior to re-transplantation (CAV #10) exhibited comparatively lower endothelial PTBP1 levels than other CAV samples (Figure 2B & SI Figure 7). This observation highlights the variability of endothelial PTBP1 expression between individual CAV patients and is consistent with the possibility that lower endothelial PTBP1 levels may reflect delayed disease progression.

**Figure 2.**
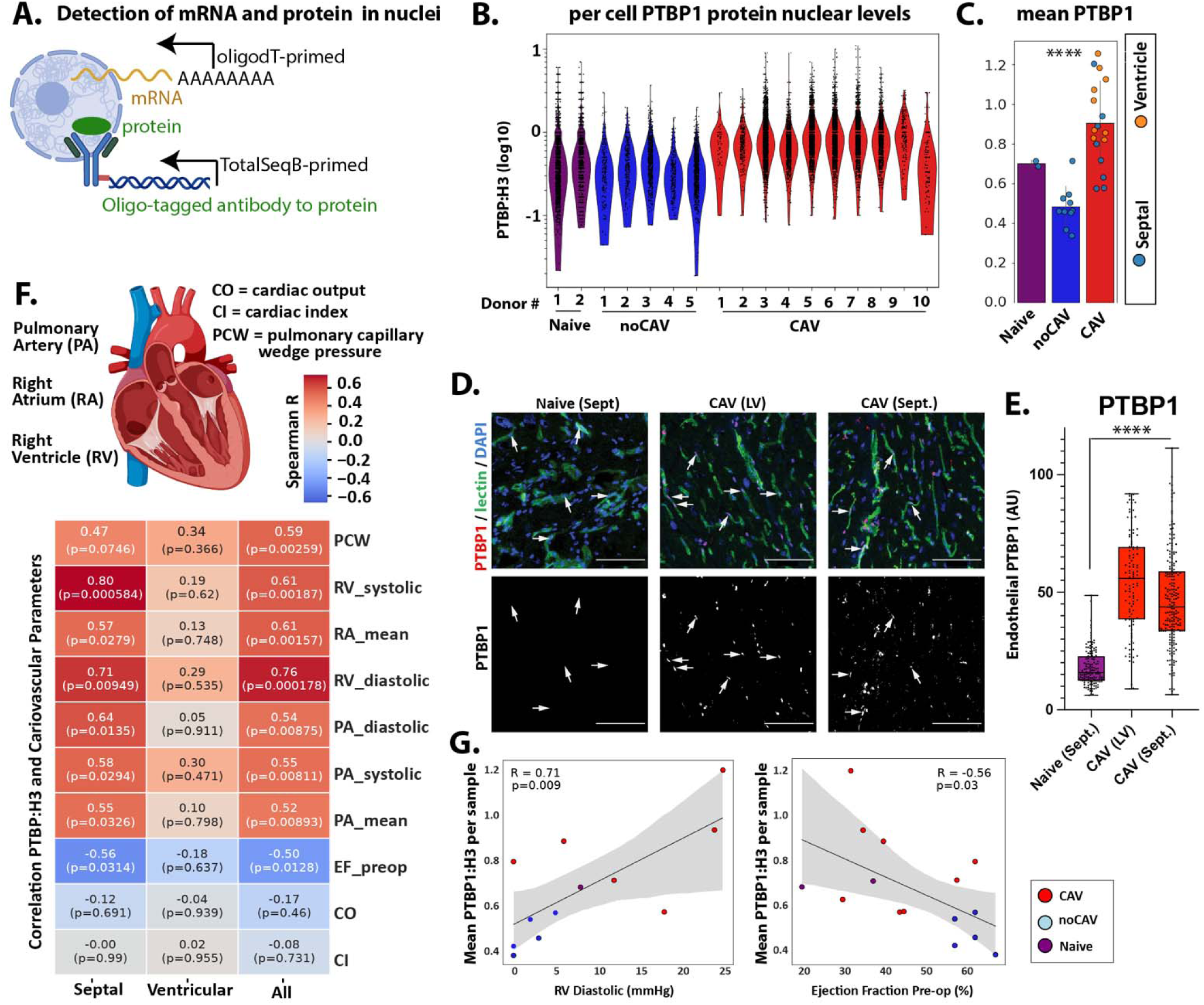
Mean endothelial nuclear levels of PTBP1 are associated with cardiac allograft vasculopathy and cardiac dysfunction. (A) inCITE-seq schematic, showing measurement of nuclear proteins (e.g. Histone H3 and PTBP1) on nuclei, along with mRNA transcripts. (B) Distribution of PTBP1 levels, normalized to Histone H3 on each septal endothelial nucleus from each donor. CAV = explant cardiac allograft vasculopathy, Biopsy = biopsy during routine examination, without CAV, and Nontransplant = heart tissue from hearts never transplanted. (C) Mean PTBP1 levels in each sample, with the inclusion of means from ventricle samples as well. Significance by one-way ANOVA, with post-hoc Tukey’s test showing difference between CAV and biopsied non-CAV samples. (D) Immunofluorescence staining, showing PTBP1 levels in DAPI+ nuclei associated with UEA-1 lectin-stained vasculature. CAV tissues are either left ventricular (LV) or septal (Sept), or from an unaffected non-transplant heart tissue. Scale bar = 100 μm (E) Box and whisker plots showing the distribution of PTBP1 staining in left ventricular (LV) and septal (Sept). tissues. Dots represent single nuclei from three random high-powered fields per sample: Naïve (n=2), CAV-LV (n=2), CAV-Sept. (n=4) (F) Pseudobulk Spearman correlations between septal samples, ventricular samples, or all samples, and cardiovascular parameters measured in hearts, when measurements are available. (G) Scatter plots showing trends and individual values for the selected correlations, with Spearman correlation R value and significance.

CAV development is associated with progressive impairment of left ventricular systolic and diastolic function^18–22^. Given the association between endothelial PTBP1 and CAV, we next asked whether PTBP1 expression correlates with clinical measures of cardiac dysfunction in this sample cohort. Cardiovascular parameters were correlated with sample-level endothelial PTBP1 expression across CAV-positive, CAV-negative, and naïve patients. Endothelial PTBP1 levels showed a strong inverse correlation with ejection fraction, indicating an association with left ventricular systolic dysfunction (Figure 2F&G). Endothelial PTBP1 levels also positively correlated with pulmonary capillary wedge pressure (PCWP) and right atrial and right ventricular pressures, consistent with diastolic dysfunction and/or secondary changes to severe systolic dysfunction (Figure 2F&G). These associations were significant in septal samples and showed similar, but non-significant, trends in ventricular tissues where sample size was limited (Figure 2F).

Although CAV is classically defined by luminal narrowing of the epicardial and intramyocardial arteries, all parts of the vasculature including capillaries and veins are affected^23^. To determine whether elevated PTBP1 expression was restricted to specific vessel subtypes, we established an arterial-venous hierarchy within the endothelial compartment using established marker transcripts (Figure 3A, SI Figure 8A&B). The highest PTBP1-expressing nuclei (top 30%) were enriched in the arteriolar-capillary endothelium (Figure 3B), which overlapped with CAV nuclei in this same population (Figure 3C). However, when PTBP1 expression was examined at the sample level, PTBP1 levels were increased in all endothelial compartments in CAV-positive samples, and the apparent enrichment in the arteriolar endothelium represented just one of the CAV samples (SI Figure 8C). Thus, PTBP1 levels were increased in the endothelium throughout the vascular hierarchy.

**Figure 3.**
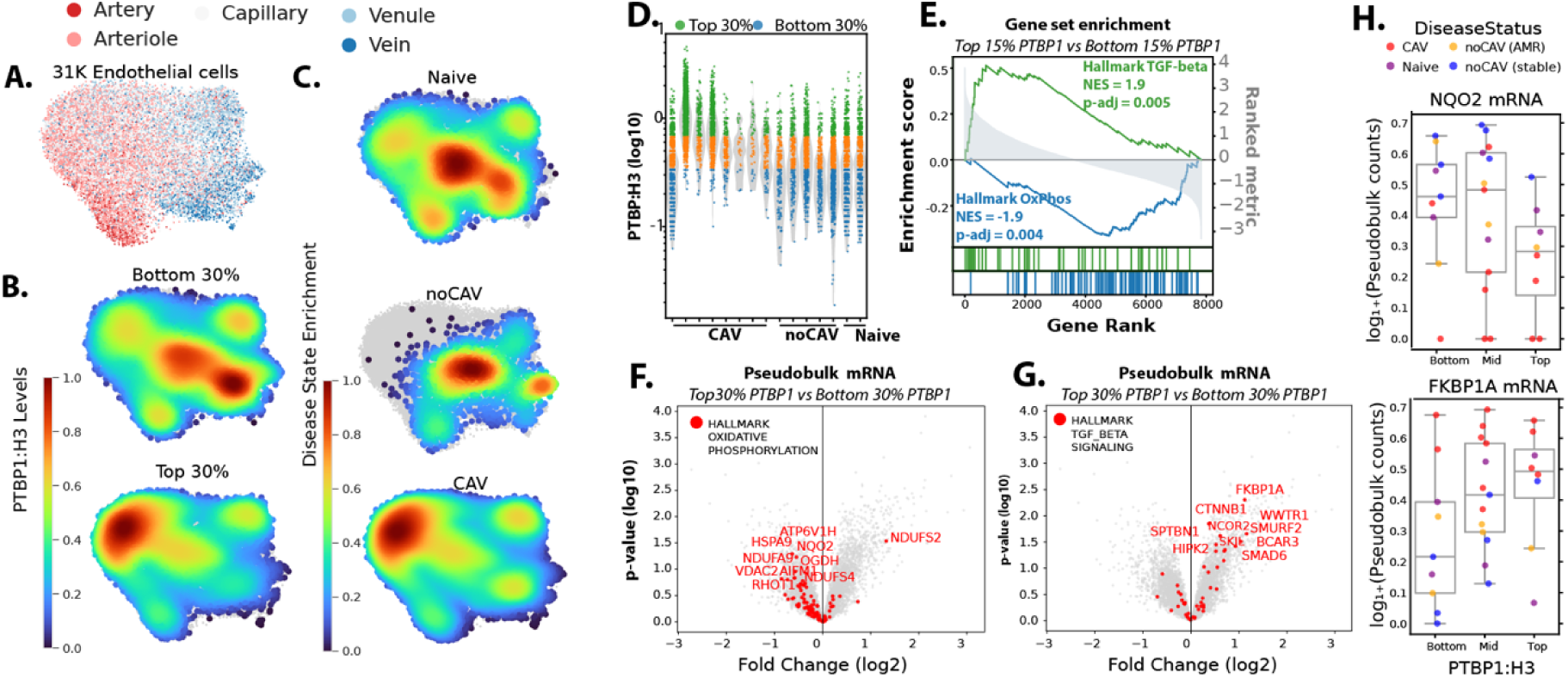
Endothelial PTBP1 levels correlate with mRNA signatures of allograft vasculopathy at a single cell level. (A) Endothelial annotated inCITE-Seq nuclei, after filtering by minimum H3 expression. (B) Location of highest density of PTBP1:H3 top 30% and bottom 30% of all nuclei. (C) Location of highest density of disease specific nuclei. (D) Violin plot showing the distribution of PTBP1, normalized to histone H3 in each nucleus, with the top and bottom 30% of all nuclei highlighted. (E) Hallmark pathway enrichment following pseudobulk analysis of the mRNA levels in the top 30% PTBP1 nuclei versus the bottom 30% nuclei, after filtering to samples with >100 endothelial nuclei passing H3 level cutoff. (F&G) Volcano plots for pseudobulk altered mRNA transcripts, indicating specific genes and their enrichment in nuclei with top 30% vs bottom 30% of PTBP1:H3 expression. (H) Plots of individual genes from OxPhos (NQO2) or TGF-β (FKBP1A) signatures, showing altered expression in cells with bottom 30% of PTBP1:H3 expression or top 30% of PTBP1:H3 expression.

Collectively, these inCITE-seq data demonstrate that PTBP1 is broadly elevated in endothelial nuclei across the CAV vasculature and is associated with clinical measures of cardiac dysfunction. These findings identify endothelial PTBP1 as a marker of endothelial dysfunction in the microvasculature and link endothelial reprogramming to cardiac dysfunction during CAV development.

### Endothelial PTBP1 expression correlates with TGF-β-driven transcriptional signatures and suppression of mitochondrial metabolic pathways

Next, we used a transcriptome-wide approach to identify signaling pathways associated with high nuclear PTBP1 expression in endothelial cells. Endothelial nuclei within each sample were stratified into PTBP1-high (top 15% expression) and PTBP1-low (bottom 15% expression) groups based on normalized nuclear PTBP1 levels (Figure 3D). Differential gene expression between PTBP1-high and PTBP1-low endothelial nuclei was assessed by pseudobulk analysis using DESeq2 across CAV-negative biopsy, CAV-positive, and naïve control septal samples (SI Table 4A-D). Gene set enrichment analysis (GSEA) was performed on the resulting differentially expressed genes to identify pathways associated with PTBP1 expression status (SI Table 5A-D).

Among the top Hallmark pathways enriched in PTBP1-high versus PTBP1-low-expressing nuclei in septal samples was TGF-β signaling, with PTBP1-high ECs showing increased expression of TGF-β-inducible genes such as *FKBP1A* (Figure 3E,G,H). Additional pathways enriched in PTBP1-high ECs include Hedgehog and WNT/β-catenin signaling, which are classically implicated in neointimal formation and proliferative smooth muscle phenotypes^24–27^ (SI Table 5). TGF-β signaling has previously been linked to impaired oxidative phosphorylation (OxPhos) and endothelial mitochondrial dysfunction^4,28–30^. Consistent with this, OxPhos was the most significantly downregulated Hallmark pathway in PTBP1-high compared with PTBP1-low endothelial nuclei (Figure 3E,F,H). These transcriptional differences were evident at both the pathway level and in individual gene expression across samples from distinct disease states (Figure 3H). In addition, metabolic pathways of fatty acid metabolism and glucose metabolism were also downregulated, in line with the role of PTBP1 as a key regulator of EC metabolism^31^ (SI Table 5A-D).

Together, these data demonstrate that endothelial PTBP1-high nuclei are characterized by enrichment of TGF-β-associated transcriptional programs coupled with suppression of metabolic pathways, including oxidative phosphorylation. Thus, endothelial PTBP1 is associated with TGF-β-linked metabolic dysfunction in the microvasculature.

### Endothelial-specific PTBP1 deletion limits CAV development in murine heart transplants

Our human data demonstrated that endothelial PTBP1 is broadly increased during CAV development, where it correlates with enhanced TGF-β signaling, suppression of OxPhos-related transcripts, and cardiac dysfunction. To determine whether endothelial PTBP1 plays a causal role in CAV pathogenesis, we examined the effects of endothelial-specific *Ptbp1* deletion in a murine model of cardiac allograft vasculopathy.

We performed heterotopic heart transplantation of C57BL/6 *Ptbp1* wild-type (*Ptbp1^f/f^*, WT) or endothelial cell-specific knockout (*Cdh5(PAC)-CreERT2; Ptbp1^f/f^*, EC-KO) donor hearts into sex-matched F1 hybrid (C57BL/6 x BALB/c) recipients. In this model, donor cardiac tissues lack host MHC molecules, triggering NK cell and CD4+ T cell-dependent immune activation that drives IFN-γ-dependent neointimal proliferation, closely recapitulating key features of human CAV pathology^32–34^. Donor mice (5 months-1 year of age) were treated with tamoxifen approximately one month prior to transplantation to induce endothelial-specific deletion of *Ptbp1.* Efficient deletion of *Ptbp1* in graft endothelial cells was confirmed by immunostaining and quantitative PCR of sorted endothelial cells isolated from transplanted hearts (SI Figure 9 A-C). Grafts were analyzed 40 days post-transplant, a time point characterized by established CAV lesions comparable to those observed in human CAV samples^32^.

Histological analysis revealed a marked reduction in CAV severity in EC-KO grafts. Blinded pathological scoring demonstrated a 53 percentage-point reduction in CAV incidence in EC-KO grafts compared with WT controls. In littermate-controlled experiments, this reduction did not reach statistical significance (P=0.12), likely reflecting limited WT sample size; however, the magnitude and direction of effect were consistent with those observed in a larger historical WT cohort in the same model (56 percentage-point reduction, P=0.003) (Figure 4A). The relative reduction was similar across sexes. There was also a trend towards reduced intimal hyperplasia in EC-KO compared with WT grafts (Figure 4B&C). Notably, the degree of protection conferred by endothelial *Ptbp1* deletion was comparable to prior studies using combined antibody-mediated depletion of CD4+ T cells, CD8+ T cells, and NK cells^32^ (Figure 4A). In addition to reduced vascular pathology, EC-KO grafts were also partially protected from post-transplant cardiac atrophy, assessed by heart weight (Figure 4D).

**Figure 4.**
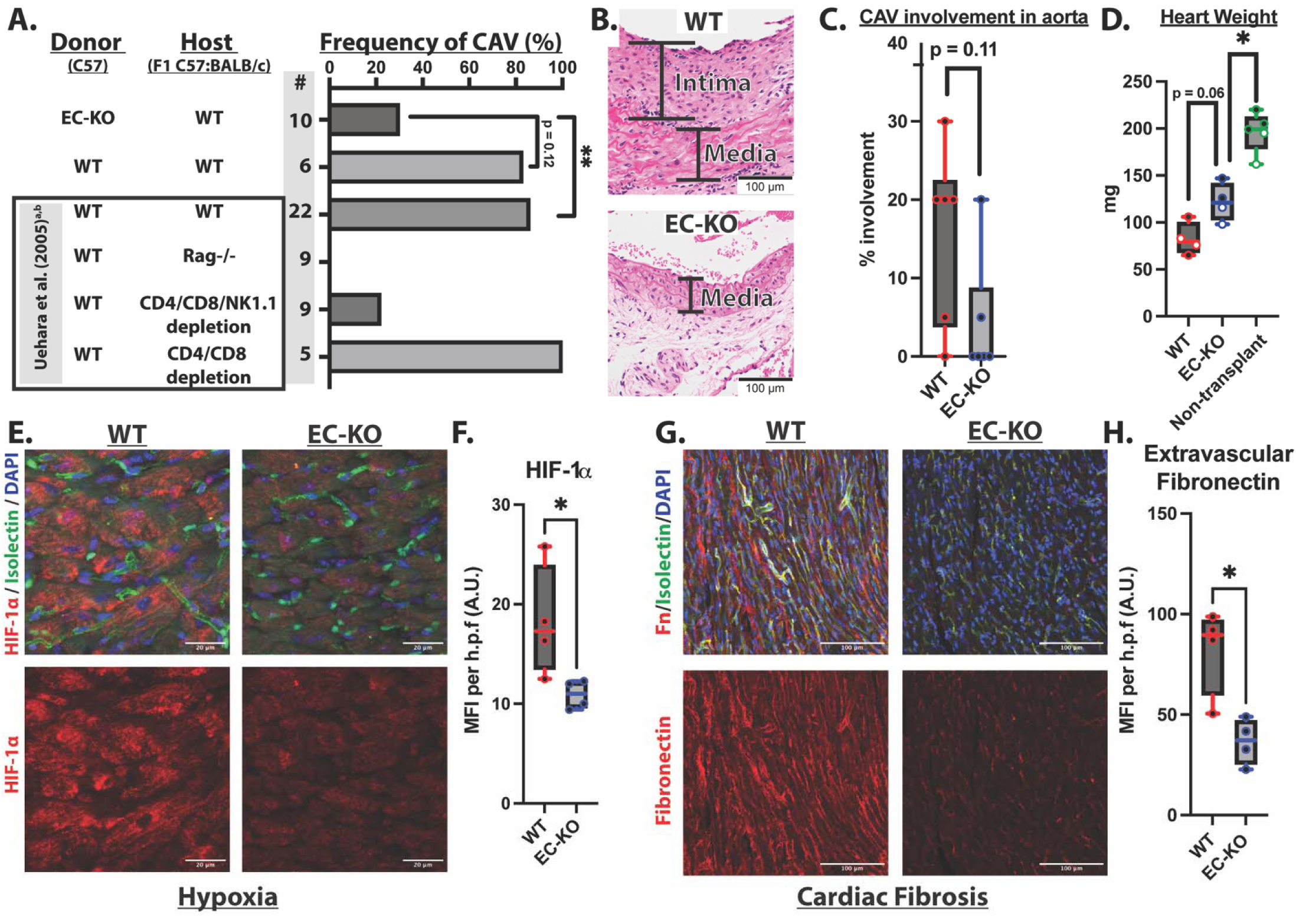
PTBP1 EC-KO reduces CAV development, cardiac fibrosis, and hypoxia in a murine F1 hybrid heart transplant model. (A) Blinded assessment of intimal proliferation in aorta of transplanted heart tissues by a trained pathologist at 40 days post-transplant, as a measure of cardiac allograft vasculopathy, with comparison to prior data in the model (Uehara *et al*.), with differences by Fisher’s exact test. EC-KO: n = 10 mice (7 male, 3 female); WT: n = 6 mice (4 male, 2 female) (B) Representative images of aortic sections stained with Hematoxylin and Eosin (H&E), and marking neointima. (C) Detailed quantification of CAV involvement in a subset of the samples (WT, n = 6 and EC-KO, n = 6) (D) Quantification of total heart weight post-transplant for WT and EC-KO grafts and non-transplant age-matched controls. White dots = female, Black dots = male mice (E) Representative immunofluorescence images of ventricular tissue from WT and EC-KO grafts stained with HIF-1α(red), isolectin B4 (green), and DAPI (blue). Composite and single-color HIF-1α staining is shown (63x). (F) Mean fluorescence intensity of HIF-1α in grafts (WT, n = 4 and EC-KO, n = 4). (G) Representative fibronectin (red), isolectin B4 (green), and DAPI (blue) staining of the same graft ventricular tissues (20x). (H) Mean fluorescence intensity of extravascular fibronectin staining quantified by masking and removing isolectin+ area (WT, n = 4 and EC-KO, n = 4).(F,H) Mean expression per mouse was calculated using three images. (C,D,F,H) Box plots show median, min, and max with mice as individual datapoints. Significance was assessed using Mann-Whitney test *P < 0.05, **P < 0.01, ***P < 0.001.

We next examined tissue-level hallmarks of CAV-associated dysfunction, including hypoxia^35–38^ and fibrosis^39,40^. Immunostaining for hypoxia-inducible factor 1α (HIF-1α) revealed an expected increase in transplanted grafts, relative to non-transplanted heart tissues, which was reduced in EC-KO grafts (Figure 4E-F, SI Figure 9D-F). Extravascular fibronectin deposition within the ventricular myocardium—a marker of fibrotic remodeling—was also reduced in EC-KO grafts compared with WT transplants, consistent with reduced fibrosis (Figure 4G-H).

Collectively, these findings demonstrate that endothelial-specific deletion of *Ptbp1* prior to heart transplantation markedly limits CAV development, preserving myocardial integrity while reducing hypoxic and fibrotic remodeling. These results establish endothelial PTBP1 as a key driver of graft-intrinsic vascular pathology in cardiac allograft vasculopathy.

### Endothelial PTBP1 deletion limits CAV-associated mitochondrial dysfunction

Endothelial cells play a central role in CAV pathogenesis through their capacity to regulate immune cell activation and were the direct target of PTBP1 deletion in our model. To define endothelial-intrinsic and immune-mediated changes contributing to reduced CAV development in EC-KO grafts, we isolated immune and non-immune cells from transplanted hearts (4 WT, 4 EC-KO samples) and performed cellular indexing of transcriptomes and epitopes by sequencing (CITE-seq). This approach enables simultaneous quantification of transcriptomic and surface protein expression across 102 murine immune-related markers^41^ (SI Table 6). After quality control and filtering, we retained ∼7,400 cells from WT grafts and ∼10,500 cells from EC-KO grafts. Joint RNA-protein-based clustering and UMAP projection identified the major cardiac cell populations (Figure 5A-B and SI Figure 10). Importantly, RNA and surface protein expression of canonical immune markers (e.g., CD4, CD8) showed strong concordance (SI Figure 10E), validating the integrated CITE-seq approach.

**Figure 5.**
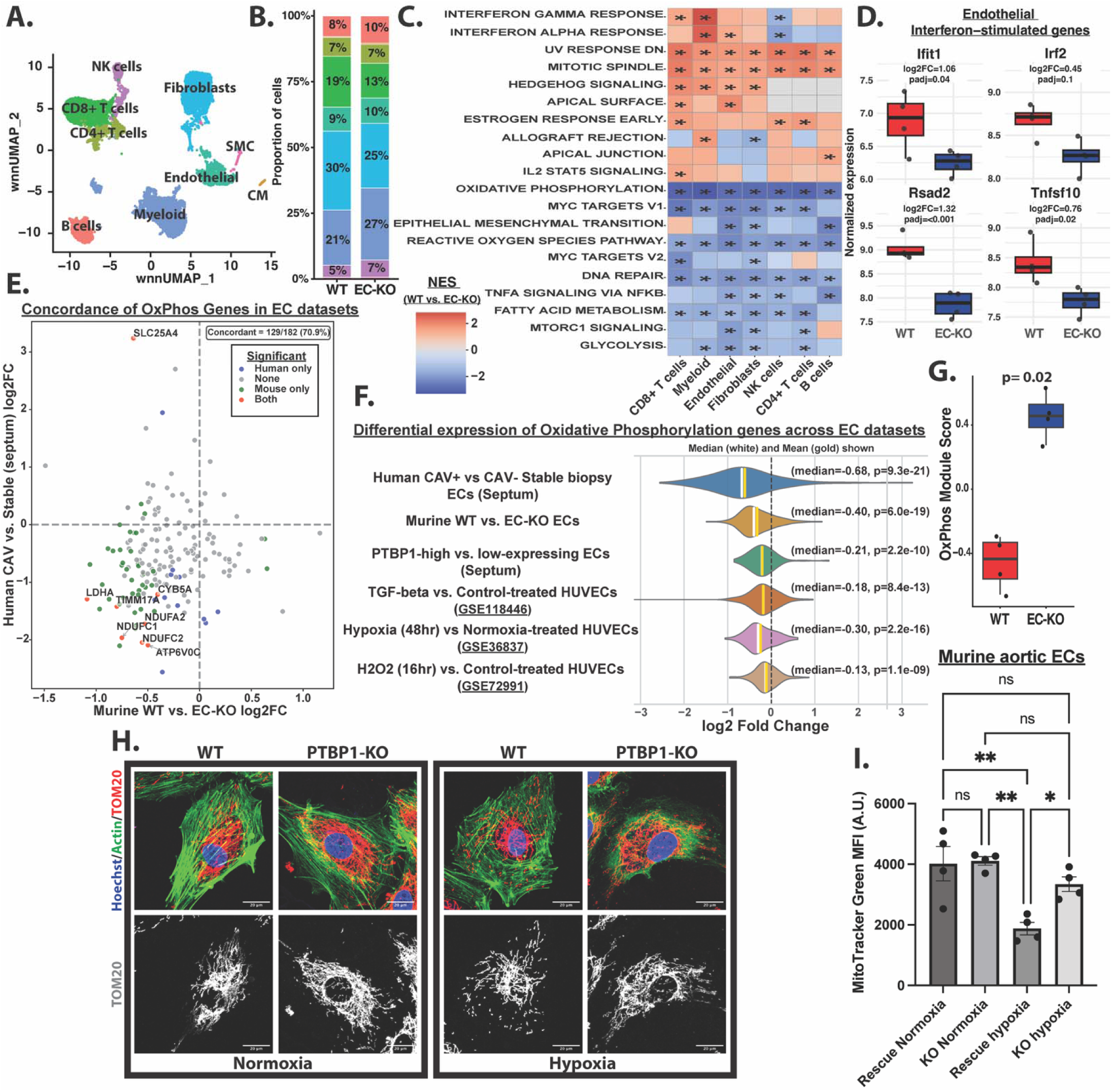
Endothelial oxidative phosphorylation is decreased in human and murine cardiac allograft vasculopathy. (A) Weighted nearest neighbor (WNN) RNA–protein integrated UMAP of cardiac cell types identified by CITE-seq. (B) Relative proportions of major cardiac cell types in WT and EC-KO grafts from CITE-seq data. (C) Heatmap showing the top 10 upregulated and downregulated Hallmark pathways across major cell types comparing WT vs. EC-KO grafts, based on gene set enrichment analysis of pseudobulk differential gene expression. NES, Normalized enrichment score; stars denote FDR-adjusted q values (q<0.1). (D). Pseudobulk expression of select interferon-stimulated genes in endothelial cells from WT and EC-KO grafts. Bar plots represent variance-stabilizing transformation (VST)-normalized expression, log2FC and adjusted p-values were calculated using DESeq2. (E) Scatter plot comparing changes in expression of oxidative phosphorylation genes (Hallmark Oxidative Phosphorylation gene set) between human (CAV+ vs. CAV- stable biopsies) and murine (WT vs EC-KO) endothelial datasets. Genes significantly differentially expressed (adjusted p < 0.1) in either dataset are labeled. Concordance of altered oxidative phosphorylation genes between species is indicated. (F) Median log2 fold-change of Hallmark Oxidative Phosphorylation genes across multiple endothelial datasets. Differential expression for public datasets was assessed using GEO2R. A binomial test was used to determine if the median log2FC was significantly different than zero (null hypothesis). (G) Pseudobulk module score of Hallmark Oxidative Phosphorylation gene set between WT and EC-KO grafts. Statistical significance was assessed using the Mann-Whitney U test (H) Representative immunofluorescence images of WT and PTBP1 KO HUVECs treated +/- hypoxia for 48 hrs and stained with TOM20 (mitochondrial marker), phalloidin (actin), and Hoechst (upper panel). Corresponding TOM20 staining (grayscale)(bottom panel). (I) Flow cytometric analysis of murine aortic endothelial cells cultured +/- hypoxia for 48 hrs and stained with MitoTracker Green. Statistical significance was determined by one-way ANOVA with Tukey’s multiple comparisons test.

To identify transcriptomic differences across endothelial and other major cell populations, we performed pseudobulk differential gene expression analysis comparing WT and EC-KO heart grafts, followed by gene set enrichment analysis (GSEA) to identify differentially regulated pathways (SI Table 7&8). Interferon signaling is a well-established contributor to CAV pathogenesis^32,42–44^, and consistent with this, WT grafts exhibited increased IFN-γ and IFN-α signaling across multiple cell types including endothelial, fibroblast, myeloid, and CD8+ T cells, relative to EC-KO grafts (Figure 5C).

Focusing specifically on endothelial cells, WT grafts showed enrichment of interferon signaling and increased expression of canonical interferon-stimulated genes, including *Ifit1*, *Rsad2*, *Irf2*, and *Tnfsf10* (Figure 5C&D, SI Figure 11A). Endothelial cells from WT grafts were also enriched for Hedgehog signaling and mitotic spindle-associated pathways (Figure 5C). In contrast, endothelial cells from EC-KO grafts were enriched for metabolic pathways including OxPhos, glycolysis, and fatty acid metabolism, as well as pathways related to reactive oxygen species (ROS) (Figure 5C). Among the ROS-associated genes upregulated in EC-KO endothelial cells were several involved in antioxidant defense, including *Gpx1*, *Prdx1*, and *Selenow* (SI Figure 11C), suggesting enhanced oxidative stress resilience. Notably, canonical TGF-β signaling did not differ significantly between WT and EC-KO grafts (SI Figure 11D), suggesting that PTBP1 likely modulates downstream consequences of pro-fibrotic signaling.

We next examined differences in surface protein expression between WT and EC-KO endothelial cells. WT endothelial cells displayed elevated expression of CD36, a scavenger receptor that facilitates fatty acid uptake and has been linked to endothelial stiffening, vascular inflammation, and atherogenesis (SI Figure 11B)^45,46^. Elevated CD36 has been associated with lipid accumulation-induced mitochondrial dysfunction^47,48^, consistent with the suppression of OxPhos observed in WT graft endothelium. Despite our prior findings that PTBP1 deletion attenuates TNF-induced adhesion molecule expression *in vitro*^11^, we did not detect significant differences in surface ICAM1 or VCAM1 expression between WT and EC-KO graft endothelium *in vivo* (SI Figure 11B)(SI Table 7). Expression of additional endothelial immune-regulatory molecules, including MHC-II, PD-L1, and CD200, was similarly unchanged between groups (SI Figure 11B). Together, these findings suggest that the protective effect of endothelial PTBP1 deletion is unlikely mediated by leukocyte recruitment or antigen presentation, but rather modulation of endothelial metabolic and inflammatory state.

To determine whether endothelial transcriptional changes observed in murine grafts mirrored those in human CAV, we compared significantly differentially expressed genes from human transplant endothelium (CAV-positive explants versus CAV-negative stable biopsies) with those from mouse heart graft endothelium (WT versus EC-KO). This analysis identified 146 shared genes, most of which (73.3%, 107 genes) were concordantly regulated—upregulated in human CAV and reversed by endothelial PTBP1 deletion in mice or downregulated in human CAV and increased in the endothelium of EC-KO murine grafts (SI Figure 11E&F, SI Table 9). This overlap was significantly greater than expected by chance (Fisher’s exact test, two-sided P < 0.0001, odds ratio = 6.2), suggesting a link between endothelial changes in human and murine CAV development. Concordantly upregulated genes were enriched for insulin signaling, whereas concordantly downregulated genes were enriched for OxPhos and aerobic respiration pathways (SI Figure 11 G&H, SI Table 10), highlighting endothelial metabolic reprogramming as a conserved feature of CAV development.

Consistent with prior reports linking CAV to mitochondrial dysfunction^49^, OxPhos-associated gene expression was markedly reduced in endothelial cells from human CAV-positive explants and WT mouse grafts (Figure 5E-G). Similar suppression of OxPhos genes was observed in endothelial cells exposed to profibrotic and inflammatory stressors relevant to CAV, including TGF-β^50^, hypoxia^51^, and oxidative stress^52^ (Figure 5F), supporting a convergent mitochondrial stress response that promotes endothelial dysfunction and immunogenicity^9,10,53^.

PTBP1 is a known regulator of cellular metabolism and mitochondrial function^54^. Given that hypoxia exacerbates endothelial mitochondrial dysfunction^55^, and is increased in both human and murine CAV grafts, we next tested whether PTBP1 contributes to hypoxia-induced endothelial mitochondrial impairment. Wild-type (WT) or PTBP1-knockout (KO) human umbilical vein endothelial cells (HUVECs) were exposed to hypoxia (1% O_2_) for 48 hours and mitochondria were assessed using TOM20 immunostaining. Under hypoxic conditions, WT endothelial cells exhibited increased mitochondrial fission, characterized by fragmented mitochondria with reduced mitochondrial density, whereas PTBP1-KO cells had preserved mitochondrial structure and content (Figure 5H).

Consistent with these findings, murine aortic endothelial cells exposed to hypoxia showed reduced mitochondrial content when PTBP1 expression was restored in PTBP1-deficient cells (Rescue), whereas mitochondrial content remained preserved in PTBP1-KO cells (Figure 5I). These data indicate that PTBP1 expression is sufficient to promote hypoxia-induced mitochondrial loss.

Excessive mitochondrial fission and mitochondrial dysfunction can promote the release of mitochondrial DNA and RNA into the cytosol, where they activate innate immune signaling pathways, leading to induction of a type I interferon response^56,57^. To assess whether PTBP1-dependent mitochondrial dysfunction was associated with interferon signaling under hypoxic stress, we stained HUVECs for IFIT1 following 48 hours of hypoxia. WT cells showed a robust increase in IFIT1 expression under hypoxic relative to normoxic conditions, whereas this response was markedly attenuated in PTBP1-KO cells (SI Figure 12B&C).

To define potential molecular mechanisms underlying this mitochondrial phenotype, we reanalyzed alternative splicing data from murine PTBP1-KO and Rescue endothelial cells^11^. This analysis identified numerous mitochondrial-associated genes that underwent PTBP1-dependent alternative splicing (ΔPSI >0.2) following PTBP1 deletion (SI Figure 12 D-F). Among these targets, Mff, a key regulator of mitochondrial fission, exhibited increased inclusion of exons generating an isoform previously associated with reduced mitochondrial fission and dampened immune responses^58^. In addition, PTBP1 deletion promoted exclusion of exons 5 and 6 of Immt, which encodes the MIC60 protein, a component of the MICOS complex involved in mitochondrial structural maintenance^59^. Loss of these exons results in removal of a disordered and coiled-coil region from the encoded protein. Notably, similar human splice isoforms of *Immt* lacking exon 6 have been associated with healthy human cardiac tissue and normal postnatal cardiac development, although its precise functional role remains undefined^60,61^.

Together, these results demonstrate that endothelial PTBP1 deletion preserves mitochondrial content and integrity under CAV-associated stress, likely through alternative splicing of key mitochondrial regulators. These findings identify PTBP1 as a central mediator of endothelial mitochondrial dysfunction and inflammatory signaling in cardiac allograft vasculopathy.

### Loss of endothelial PTBP1 attenuates T and NK cell activation in murine cardiac grafts

Because mitochondrial dysfunction and interferon signaling can enhance EC immunogenicity within heart transplants^10,53,62^, we asked whether endothelial PTBP1 deletion alters graft immune phenotypes. In the parental-to-F1 hybrid murine CAV model, NK cells and T lymphocytes are critical mediators of disease, as their combined depletion markedly limits CAV development^32^ (Figure 4A).

Based on our prior findings that PTBP1 promotes endothelial adhesion molecule expression and myeloid accumulation in atherosclerotic plaques^11^, we investigated whether endothelial PTBP1 deletion altered immune cell recruitment in cardiac grafts. Total immune cell numbers were similar in WT and EC-KO grafts at the experimental endpoint and were increased over naïve heart tissues (Figure 6A). This lack of difference in immune infiltration is consistent with the similar endothelial adhesion molecule expression observed between WT and EC-KO grafts (SI Figure 11B). Moreover, the composition of T, NK, and B cell populations in recipient spleens was unchanged between groups, suggesting minimal effects on systemic immunity (SI Figure 13A).

**Figure 6.**
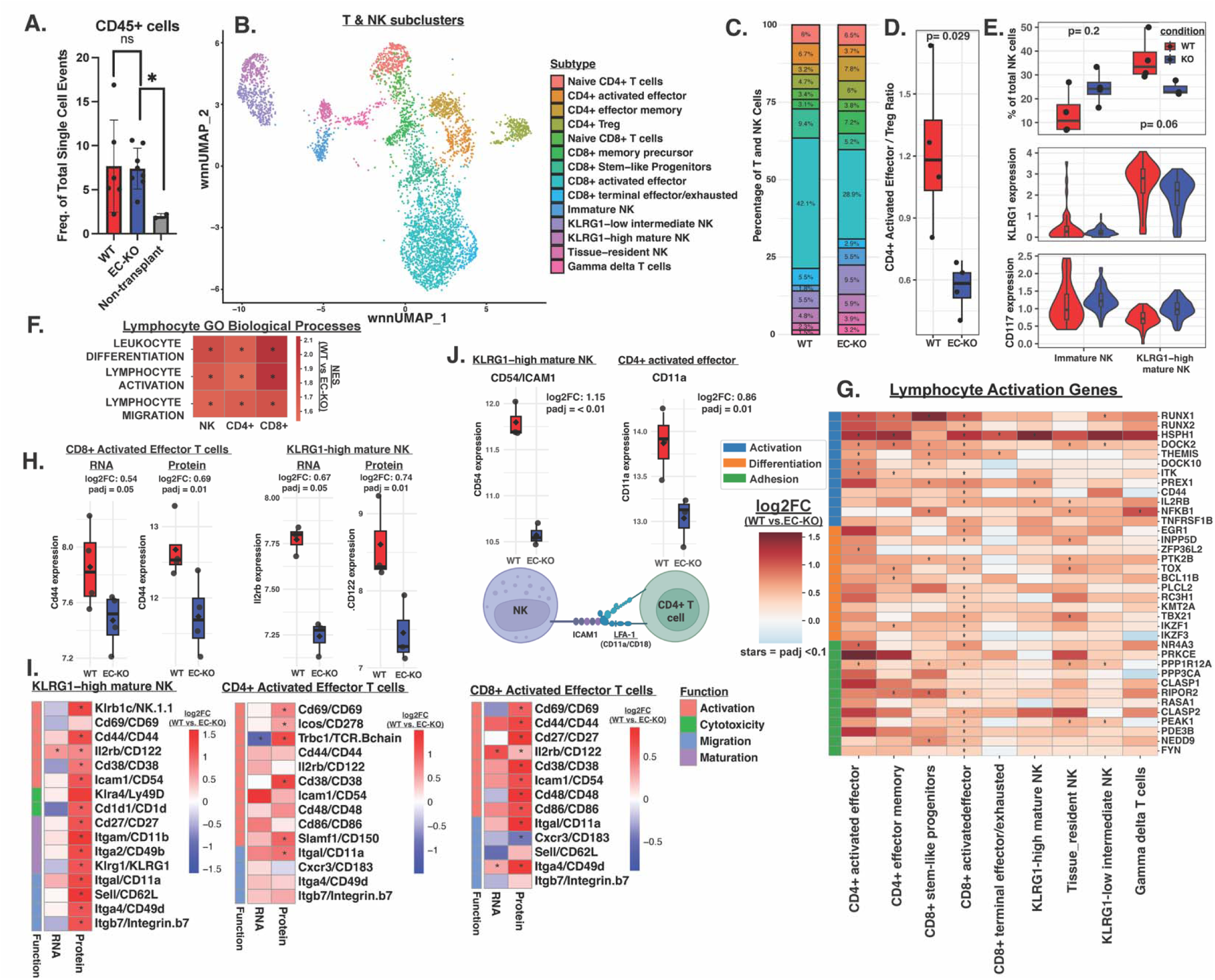
T and NK lymphocytes from CAV-prone WT transplants exhibit increased markers of activation, differentiation, and adhesion compared to EC-KO transplants. (A) Flow cytometry analysis of CD45+ cells isolated from transplanted hearts (WT, n = 4; EC-KO, n = 4) and non-transplant controls (n = 2). Data are shown as mean ± SD. Statistical significance was assessed using Mann-Whitney U test (B) Weighted nearest neighbor (WNN) UMAP of T and NK lymphocytes from WT and EC-KO cardiac transplants (n = 4 per condition), colored by subtype. (C) Stacked bar plot showing proportions of T and NK lymphocyte subtypes by condition. (D) Ratio of CD4+ activated effector T cells to CD4+ Treg cells per sample. (E) NK cell maturation in WT versus EC-KO grafts. Top: Proportion of immature and KLRG1-high mature NK cells among total NK cells per sample. Middle: Centered log-ratio (CLR)–normalized KLRG1 protein expression by NK subtype. Bottom: CLR-normalized CD117 (c-KIT) protein expression by NK subtype. (F) GSEA of ranked pseudobulk differentially expressed genes (WT vs. EC-KO) in broad T and NK lymphocyte populations, showing NES for representative Gene Ontology Biological Process (GOBP) pathways (* FDR q < 0.1). (G) Heatmap of differentially expressed lymphocyte activation–related genes across T and NK subtypes (WT vs. EC-KO) by pseudobulk analysis (n = 4 per condition). Genes were grouped by Gene Ontology category; genes present in multiple categories were assigned hierarchically (Activation > Differentiation > Adhesion) (* padj < 0.1). (H) Pseudobulk RNA and protein expression of Cd44/CD44 in CD8+ activated effector T cells (left) and Il2rb/CD122 in KLRG1-high mature NK cells (right). Dots represent individual samples. Line indicates median; diamond indicates mean. For EC-KO KLRG1-high mature NK cells n =3. (I) Heatmap of differentially expressed RNA and protein markers of lymphocyte function in effector T and NK subtypes (WT vs. EC-KO) by pseudobulk analysis (* padj < 0.05). (J) CLR-normalized pseudobulk expression of CD54 (ICAM-1) in KLRG1-high mature NK cells and CD11a (LFA-1) in CD4+ activated effector T cells (WT vs. EC-KO). Schematic of ICAM-1–LFA-1 interaction was created with BioRender.com. (D&E) Differences in proportions between groups were assessed using two-tailed Wilcoxon rank-sum test. (H-J) Significant differences in pseudobulk RNA and protein levels were assessed using DESeq2 which obtains raw p-values from the Wald test that are adjusted for multiple testing using the Benjamini–Hochberg (BH) false discovery rate (FDR) correction.

Accumulating evidence suggests that inhibition of endothelial mitochondrial fission reduces alloimmune responses and vascular inflammation^53,62,63^. Therefore, despite equal numbers of recruited immune cells, PTBP1 deletion may skew the immune response to a less active state. To investigate this possibility, we analyzed WT and EC-KO heart transplants using CITE-seq as described above, focusing on the T and NK cells linked to CAV. Using integrated RNA and surface protein information, we identified multiple heterogeneous CD4+ and CD8+ T cell and NK cell populations, spanning naïve, effector, regulatory, and varying maturation states (Figure 6B, SI Figure 13B & 14). Compositional analysis revealed that WT grafts were enriched for activated and progenitor CD4+ and CD8+ T cell subsets, whereas EC-KO grafts were enriched for immune-regulatory and less differentiated lymphocyte populations, including CD4+ Tregs, CD4+ effector memory T cells, Naïve CD8+ T cells, and NK cell populations (Figure 6C). Consistent with a shift toward a more regulatory CD4+ T cell landscape, EC-KO grafts exhibited a significantly reduced ratio of activated CD4+ effector T cells to CD4+ Tregs (Figure 6D). Within the NK cell compartment, endothelial PTBP1 deletion altered NK maturation states. EC-KO grafts showed a trend towards an increased proportion of Immature NK cells and a corresponding reduction in KLRG1-high mature NK cells, accompanied by increased expression of progenitor associated markers (c-KIT)^64^, and diminished terminal maturation markers (KLRG1) (Figure 6E).

To determine whether these compositional shifts were accompanied by functional differences, we performed integrated RNA and surface protein analysis of T and NK lymphocyte subsets. Pseudobulk analysis comparing WT and EC-KO grafts revealed increased expression of key lymphocyte activation markers in WT grafts, including CD44 in activated CD8+ T cells and IL2RB in mature NK cells, at both the transcript and surface protein levels (Figure 6H). Gene set enrichment analysis further demonstrated significant enrichment of pathways associated with lymphocyte activation, differentiation, and adhesion in WT relative to EC-KO grafts across multiple T and NK lymphocyte populations (Figure 6F-G, SI Figure 12A, SI Table 11&12). Consistent with these findings, activated CD4+ and CD8+ effector T cells and KLRG1-high mature NK cells in WT grafts exhibited increased surface expression of markers associated with activation, cytotoxicity, maturation, and migration, compared with EC-KO grafts (Figure 6I&J, SI Table 13). These differences were most pronounced at the protein level, highlighting the utility of using the CITE-seq approach. Given evidence that NK cells can amplify T cell activation through contact-dependent mechanisms, we examined the surface expression of the adhesion molecules ICAM-1 and LFA-1 in NK and T cells^32,65^. WT grafts showed increased expression of ICAM1 on KLRG1-high mature NK cells and LFA-1 subunit CD11a on CD4+ activated effector T cells, consistent with molecular features associated with enhanced NK-T cell interaction (Figure 6J).

Together, these data indicate that endothelial PTBP1 deletion does not reduce immune cell recruitment but instead restrains T and NK cell activation and maturation, shifting the graft immune environment to a less inflammatory and more regulatory state.

## Discussion

The donor vascular endothelium is the first cell layer encountered by the recipient immune system and is uniquely positioned to shape local immune activation and chronic vascular remodeling. RNA-binding proteins (RBPs) such as PTBP1 are key post-transcriptional regulators of endothelial responses but are frequently subjected to autoregulatory control, resulting in discordance between mRNA and protein abundance. Using inCITE-seq to quantify nuclear PTBP1 protein alongside endothelial transcriptomes in human transplant tissues, we found that endothelial PTBP1 levels were elevated across the CAV vasculature and associated with increased TGF-β signaling, suppression of OxPhos transcripts, and cardiac dysfunction.

These human data establish strong associations between PTBP1 protein levels and endothelial reprogramming. Complementary genetic studies in a murine model of CAV demonstrate that endothelial-specific deletion of *Ptbp1* significantly limited CAV development, characterized by reduced neointimal formation, hypoxia, fibrosis, and local lymphocyte activation. Molecularly, *Ptbp1* deletion prevented OxPhos-related transcriptional changes associated with metabolic dysfunction in murine grafts and preserved endothelial mitochondrial integrity under hypoxic stress, dampening interferon responses. Notably, endothelial *Ptbp1* deletion achieved a similar level of protection as antibody-mediated depletion of all CD8+ T cells, CD4+ T cells, and NK cells in the same model, suggesting that endothelial post-transcriptional regulation may be harnessed to locally alter pathogenic immune responses while sparing systemic immunity.

We propose that pro-fibrotic and metabolic stressors (including TGF- β and hypoxia) promote endothelial PTBP1 induction and/or nuclear accumulation, which in turn promotes mitochondrial remodeling and metabolic dysfunction, amplifying innate interferon signaling and downstream lymphocyte activation that contributes to CAV progression. Below, we discuss evidence supporting this model and its implications for chronic rejection.

### PTBP1 in endothelial reprogramming in vasculopathy

InCITE-seq analysis provides a powerful approach to examine the correlations between nuclear protein levels and gene expression at single-nucleus resolution. In human CAV, nuclear PTBP1 levels were strongly associated with TGF-β-related signaling. Elevated TGF-β signaling in early heart transplant biopsies correlates with later CAV development^66^, as do markers of platelet activation and microvascular coagulation^67^. Notably, platelets are a primary source of TGF-β signaling in tissue injury and can promote endothelial mitochondrial dysfunction ^68,69^. Our prior work demonstrated that PTBP1 mRNA is induced in a platelet-dependent manner in endothelial cells exposed to disturbed flow^70^, raising the possibility that platelet-derived or injury-associated signals contribute to PTBP1 induction in the transplant setting. In addition to direct induction by TGF-β^71^, downstream consequences of TGF-β signaling—including fibronectin deposition and extracellular matrix stiffening—may further enhance PTBP1 nuclear localization, as PTBP1 functions as a mechanosensitive splice factor^72^.

Importantly, although PTBP1 levels were associated with TGF-β-related signaling in human tissues, endothelial-specific deletion of *Ptbp*1 in murine grafts did not broadly alter canonical TGF-β pathway transcripts, aside from genes involved in metabolic programming. These findings suggest that PTBP1 does not function as a primary regulator of upstream TGF-β signaling, but instead modulates selective downstream or parallel pathways, particularly those involved in metabolism. In this framework, inflammatory and pro-fibrotic cues such as TGF-β may induce PTBP1 expression, while PTBP1 shapes the metabolic consequences of these signals within endothelial cells. Consistent with this model, OxPhos and related metabolic pathways were no longer suppressed in EC-KO murine grafts, as they were in PTBP1-high endothelial nuclei in human CAV samples. Together, these findings position PTBP1 as a mediator linking TGF-β signaling and fibrotic pathways to endothelial metabolic remodeling.

### Metabolic reprogramming of the endothelium and vasculopathy

Pro-inflammatory induction of adhesion molecules and HLA expression is often viewed as the principal mechanism by which endothelial cells promote graft inflammation. However, the metabolic consequences of inflammatory signaling—and how endothelial metabolic state shapes immune responses—have received comparatively less attention. Endothelial metabolic state can influence immune function through stress signaling pathways and metabolic crosstalk with infiltrating leukocytes^56,73–76^. Based on our prior findings that PTBP1 deletion impairs TNF-induced adhesion molecule expression^11^, we anticipated that EC-KO grafts would exhibit reduced immune cell recruitment. Instead, endothelial adhesion molecule expression and total immune cell numbers in grafts were unchanged, whereas lymphocyte activation and maturation were selectively attenuated in EC-KO grafts. Restoration of OxPhos-related transcripts in EC-KO endothelium coincided with this reduced lymphocyte activation, suggesting that preventing mitochondrial dysfunction may limit alloimmune responses.

Cardiac allograft vasculopathy is characterized by diminished mitochondrial oxidative function in human cardiac grafts^49^, and multiple CAV-relevant stressors—including TGF-β, hypoxia, and oxidative stress—suppress OxPhos-related transcripts in endothelial cells^50–52,77^. These cues also promote mitochondrial fission, a process linked to enhanced innate immune activation and increased endothelial immunogenicity^10,29,53,57,62,69^. Consistent with this paradigm, PTBP1 has previously been shown to shift endothelial metabolism toward glycolysis at the expense of oxidative phosphorylation, and PTBP1 knockdown rescued mitochondrial oxidative function^14^. *In vivo*, EC-KO grafts exhibited reduced type I and type II interferon signaling. *In vitro*, PTBP1 deletion in HUVECs attenuated hypoxia-induced mitochondrial fission and IFIT1 induction. Notably, STING pathway activation – which can drive interferon type I signaling in response to mitochondrial DNA release—is required for cardiomyopathy in response to pressure overload^78^. Together, these findings support a model in which PTBP1 contributes to stress-induced mitochondrial remodeling that amplifies innate immune signaling – perhaps through STING activation, whereas PTBP1 deficiency preserves mitochondrial integrity and dampens inflammatory activation.

### Microvascular injury in chronic allograft vasculopathy

CAV is classically defined by intimal proliferation of epicardial coronary arteries. However, impaired myocardial blood reserve—a measure of microvascular function—predicts adverse outcomes after heart transplantation, even in the absence of overt epicardial stenosis^79^, indicating that microvascular dysfunction can independently contribute to graft failure. Because routine surveillance biopsies are obtained from septal tissue, our human analysis focuses on the microvasculature. Despite this constraint, we observed robust changes in septal endothelial PTBP1 protein expression and TGF-β-associated transcripts that correlated with clinical indices of systolic and diastolic dysfunction, including reduced ejection fraction and elevated diastolic filling pressures. These findings suggest that endothelial reprogramming within distal microvascular beds may contribute meaningfully to cardiac dysfunction in CAV.

Although we cannot exclude the possibility that microvascular changes in human samples occur secondary to upstream epicardial disease, our murine data support a broader role for endothelial PTBP1 across vessel sizes. Endothelial-specific *Ptbp1* deletion reduced neointimal formation in larger arteries and attenuated hypoxia and fibrosis within distal vascular beds, indicating that endothelial PTBP1 influences both epicardial remodeling and downstream microvascular injury.

In summary, these findings identify PTBP1 as a key endothelial regulator of inflammatory and metabolic reprogramming in chronic cardiac allograft rejection. Our results reveal post-transcriptional control of endothelial metabolism as a previously unrecognized mechanism contributing to CAV development and nominate the targeting of endothelial PTBP1 as a potential therapeutic approach to limit chronic graft injury while minimizing systemic immunosuppression.

## Material and Methods

An Extended Methods section is provided in the Supplementary Material.

### Human inCITE-seq analysis

#### Nuclei isolation and sequencing

De-identified heart tissues were obtained from frozen tissue remnants collected after endomyocardial biopsy, CAV-positive explant, severe rejection explant, or from non-transplanted hearts. Biopsy, CAV-positive explant, and non-transplanted samples were acquired from Dr. Nicole Valenzuela and Dr. Jeffrey Hsu (UCLA) under IRB21-1330 and IRB22-0659. Additional CAV-positive explants and severe rejection tissues were acquired from Dr. Dawn Bowles (Duke) under IRBPro00005621. All human tissue collection and analysis were performed in accordance with institutional guidelines and approved IRB protocols.

Single-nucleus inCITE-seq was performed largely as described previously (Omar et al 2025^15^), with minor modifications optimized for low-input biopsy samples and incorporation of a newly generated antibody panel. Antibodies were conjugated to DNA oligonucleotides using click chemistry, cleaned using size exclusion chromatography, pooled into a single panel, and validated prior to use. For this study, analyses focused on PTBP1, Histone-H3, and non-specific IgG controls. Detailed antibody-oligonucleotide conjugation protocols are provided in the Extended Methods.

Frozen tissue was homogenized and nuclei were isolated using Nuclei EZ Lysis Buffer. Nuclei from endomyocardial biopsy samples were labeled with distinct oligonucleotide hashtags during isolation. Following filtration and washing, nuclei were incubated with DAPI, the pooled inCITE-seq antibody panel, and Nuclear Pore Complex TotalSeq-B hashing antibodies after blocking with human Fc block and single-stranded oligonucleotides.

For ventricular samples, endothelial nuclei were additionally labeled with ERG antibody (Alexa Fluor 647, Clone EPR3864, Abcam; 1:100) to enable enrichment. Following staining, singlet nuclei were sorted and ERG+ and ERG- nuclei were pooled at a 1:1 ratio to achieve approximately 50% endothelial representation per sample prior to sequencing. Between four and seven samples were combined per batch and processed on a 10x Chromium or Chromium X(i) platform using the 10x Genomics 3’ RNA-seq and Feature Barcode kit. Libraries were prepared as previously described^15^, with a modified extension step to ensure full-length coverage. Samples were sequenced to a depth of ∼100,000 reads per nucleus for gene expression, and ∼20,000 reads per nucleus for inCITE-seq and hashing barcode features.

#### Data processing

Sequencing reads were aligned to the human genome and feature barcode libraries using Cell Ranger (v8.0.1). Ambient RNA contamination was removed using CellBender (version 0.2.2). Processed count matrices were analyzed in Jupyter Lab, following workflows adapted from Omar *et al*.^15^. Sample demultiplexing was performed using Demuxem and aided by Souporcell, which leveraged expressed genetic variation to distinguish donor- and host-derived nuclei. Graft identity was assigned based on cardiomyocyte genotype, as cardiomyocytes are donor-derived and not replaced by host cells. Doublets were identified and removed using both Souporcell and Scrublet. The resulting datasets were used for downstream analysis. Full analysis scripts and notebooks are available at https://github.com/pamurphyUCONN/2026_Pathoulas

### Murine heterotopic heart transplantation model

#### Transplantation

For endothelial-specific deletion of Ptbp1, *Cdh5(PAC)-CreERT2* mice were crossed with *Ptbp1^flox/flox^* C57BL/6J mice to generate endothelial-specific knockout (EC-KO) mice [*Cdh5(PAC)-CreERT2; Ptbp1^flox/flox^*] and floxed littermate controls^80^. Mice aged 5 months-1 year of both sexes were used for experiments. Tamoxifen (Sigma) was administered intraperitoneally (1 mg per dose, once daily for three consecutive days) approximately one month prior to transplantation. To control for potential Cre-dependent effects independent of *Ptbp1* deletion, a cohort of [*Cdh5(PAC)-CreERT2; Ptbp1^flox/flox^*] mice were treated with tamoxifen or vehicle before transplantation (see Extended Methods).

Donor hearts were transplanted into F1 BALB/c:C57BL/6J recipients using a heterotopic abdominal approach, as previously described^32,81^. The donor aorta anastomosed to the recipient abdominal aorta, and the donor pulmonary artery was connected to the recipient vena cava, allowing perfusion of epicardial vessels under arterial pressure. Cardiac perfusion was confirmed intraoperatively, and graft palpation was performed twice weekly.

At 40 days post-transplantation, grafts were harvested and either fixed for histological assessment of cardiac allograft vasculopathy (CAV) or processed for single-cell analyses and immunostaining. All mice were housed and handled in accordance with protocols approved by the University of Connecticut Health Center for Comparative Medicine and Harvard Medical School and Massachusetts General Hospital.

#### Histological assessment of cardiac allograft vasculopathy

Explanted murine hearts were fixed in 10% buffered formalin and processed for routine histology. The hearts were embedded at their base and serial sections of the aorta and coronary arteries were taken. Representative and serial sections along the ventricles were also taken. Assessment of CAV was performed by a pathologist (IR) blinded to sample identities. CAV was evaluated along the aorta, coronary and intramyocardial arteries.

#### Immunostaining

Murine heart tissues were weighed, bisected, fixed in 4% paraformaldehyde, paraffin embedded, and sectioned for immunostaining. Sections were deparaffinized, antigen retrieval was performed as needed, and samples were blocked and permeabilized prior to incubation with primary antibodies or lectins. Secondary staining was performed using biotin-conjugated antibodies against primary antibody host species. Detection was performed using streptavidin-conjugated tertiary fluorescent antibodies, and nuclei were counterstained with DAPI. Imaging was performed on a Zeiss LSM 880 confocal microscope using identical acquisition parameters across samples. Quantitation was performed using raw .CZI files in ImageJ.

Human tissues were embedded in OCT, sectioned, briefly fixed, stained, and analyzed using analogous procedures. Sudan black was used to reduce autofluorescence prior to immunostaining.

### Murine CITE-seq and flow analysis

#### Cell isolation for CITE-seq and flow cytometry analysis

Murine heart tissues were enzymatically digested to generate single-cell suspensions optimized for preservation of surface antigens. Cells were filtered, red blood cells were lysed, and viable cells were counted prior to downstream analysis. For CITE-seq, 500,000 cells per sample were used for oligonucleotide-conjugated antibody staining. Remaining cells were used for traditional flow cytometry analysis.

#### CITE-seq analysis of immune cell composition

Single-cell suspensions were blocked with murine Fc block and stained with the Mouse TotalSeq-B Universal Cocktail (BioLegend) and TotalSeq-B cell hashtags (BioLegend) alongside fluorophore-conjugated antibodies used for sorting. Live cells were isolated by flow cytometry and sorted into endothelial (CD31+/ICAM2+), leukocyte (CD45+), and non-endothelial/non-immune populations. Sorted cells were pooled by genotype and processed for CITE-seq using 10x Genomics 3’ RNA-seq and Feature Barcode platform. Libraries were prepared as described previously^15^, with a 3-minute extension to provide full-length library coverage. Samples were sequenced to a depth of ∼100,000 reads per cell for gene expression, and ∼20,000 reads per cell for CITE-seq and hashing barcode features.

#### Flow-cytometry analysis

##### Heart

Immune cell composition was additionally assessed by flow cytometry in WT (n=6), EC-KO (n=8), and non-transplanted control hearts (n=2). Cells were blocked with murine Fc block, then stained with antibodies against CD45 and CD31 and viability dye prior to analysis. Data were acquired on an S6 SE sorter and analyzed using FlowJo.

##### Spleen

Immune cell composition of spleens was assessed by flow cytometry at 40 days post-transplant in WT (n=2) and EC-KO (n=4) mice. Cells were blocked with murine Fc block, then stained with antibodies against CD45, CD3, CD4, CD8, CD19, and NK1.1 and viability dye prior to analysis. Data were acquired on an S6 SE sorter and analyzed using FlowJo.

#### Single-cell RNA-seq and CITE-seq analysis

Sequencing reads were aligned to the murine genome and feature barcode libraries using Cell Ranger (v8.0.1). Ambient RNA contamination was removed using CellBender (0.2.2). Processed gene expression and antibody-derived tag (NEAT) count matrices were processed using Seurat. Cells were filtered based on detected gene counts (200–8000). Sample demultiplexing and initial doublet identification were performed using MULTIseqDemux. Additional doublets were identified and removed using scDblFinder. RNA data were normalized using SCTransform with regression of mitochondrial content and batch effects, followed by principal component analysis and Harmony-based integration. Surface protein data were normalized using centered log-ratio transformation and analyzed in parallel. Multimodal integration was performed using weighted nearest neighbor analysis, followed by UMAP embedding and graph-based clustering. Cell types were annotated based on canonical RNA and protein markers, and low-quality or contaminating clusters were excluded.

Pseudobulk differential expression analyses were performed on RNA and/or protein (NEAT) assays by aggregating cells by biological sample and cell type (minimum 10 cells per group). Differential expression was assessed using DESeq2, and pathway enrichment analysis was performed using preranked gene set enrichment analysis against MSigDB Hallmark and Gene Ontology gene sets (v2025.1), retaining pathways with FDR < 0.1.

### Endothelial cell culture and hypoxia experiments

Human umbilical vein endothelial cells (HUVECs) and mouse aortic endothelial cells were used for *in vitro* experiments. HUVECs were cultured in endothelial growth medium and used at passages <20. PTBP1-knockout (KO) HUVECs were generated using ribonucleoprotein-based CRISPR-Cas9 electroporation, with GFP-targeting guides used as non-targeting controls. PTBP1 loss was confirmed by immunoblotting, and pooled edited populations were used for all experiments.

PTBP1-Rescue and PTBP1-KO mAECs were generated as previously described^11^ and cultured in the absence of doxycycline for 48 hours prior to experimentation. For hypoxia experiments, endothelial cells were exposed to 1% O_2_ for 48 hours under otherwise standard culture conditions.

Mitochondrial content in mAECs was assessed by flow cytometry following MitoTracker Green staining. Mitochondrial structure and interferon response in HUVECs after hypoxia exposure was assessed by immunofluorescence. Cells were fixed and stained for TOM20 or IFIT1 and images were acquired by confocal microscopy. Detailed protocols for endothelial cell culture, CRISPR-Cas9 editing, hypoxia treatment, flow cytometry, and immunofluorescence are provided in the Extended Methods.

### Splicing analysis of published datasets

Alternative splicing analysis was performed using Leafcutter on previously published murine aortic endothelial cell RNA-seq data (PRJNA1142064)^82^. Splicing events were visualized using Leafviz.

### Statistical Analysis

Except for single-cell analyses, statistical testing was performed using GraphPad Prism. Two-group comparisons were conducted using two-tailed Student’s t tests or Mann-Whitney U tests, as appropriate. Comparisons involving multiple groups were analyzed using ANOVA with Sidak’s post hoc test or Kruskal-Wallis with Dunn’s correction. Biological replicates are defined as individual animals or human donors unless otherwise indicated. Statistical tests and sample sizes are indicated in figure legends.

## Data availability

Murine CITE-seq data are deposited in the SRA (PRJNA1405136). Human inCITE-seq data are undergoing dbGap data deposition to be available at the time of publication. Access to count tables (h5ad format) for murine and human single cell expression will be available through CZ CellxGene platform.

## Acknowledgements

In the course of this work, we received valuable input from the ImmunoCardiovascular Group at UConn Health. This work was supported by UConn Health startup funds from the School of Medicine and Department of Cell Biology, Center for Vascular Biology and Calhoun Cardiology Center, American Heart Association Innovative Project Award 19IPLOI34770151 (to P.A.M.); NIH National Heart, Lung, and Blood Institute Grants K99/R00-HL125727 and R01-HL150362 (to P.A.M); Marlene L. Cohen and Jerome H. Fleisch Chair (to P.A.M.) and NIH National Heart, Lung, and Blood Institute Grants L30-HL159824 and K08-HL151961 (to J.J.H.), American Heart Association Postdoctoral Fellowship 23POST916868 and TSF/STSA Resident Research Award (to K.D.), and Boehringer Ingelheim endowed chair (to A.T.V.).

---

**SI Figure 1.**
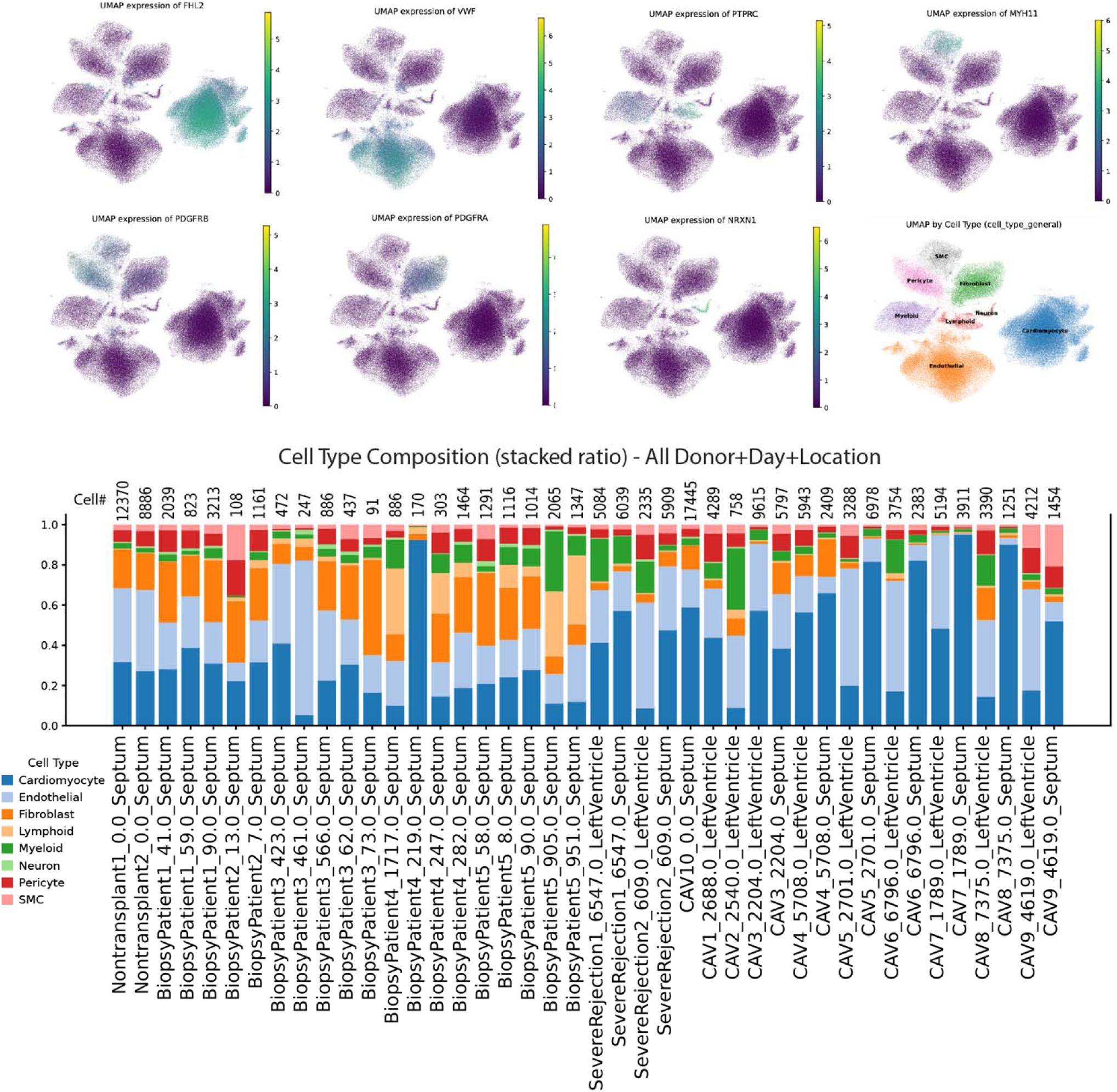
Single cell analysis showing basic characterization. UMAP clusters showing the expression of key cluster marker genes, across all samples. Barplots below show individual samples, labeled by sample ID, day post explant, and location. Total cell number in each sample is shown at the top.

**SI Figure 2.**
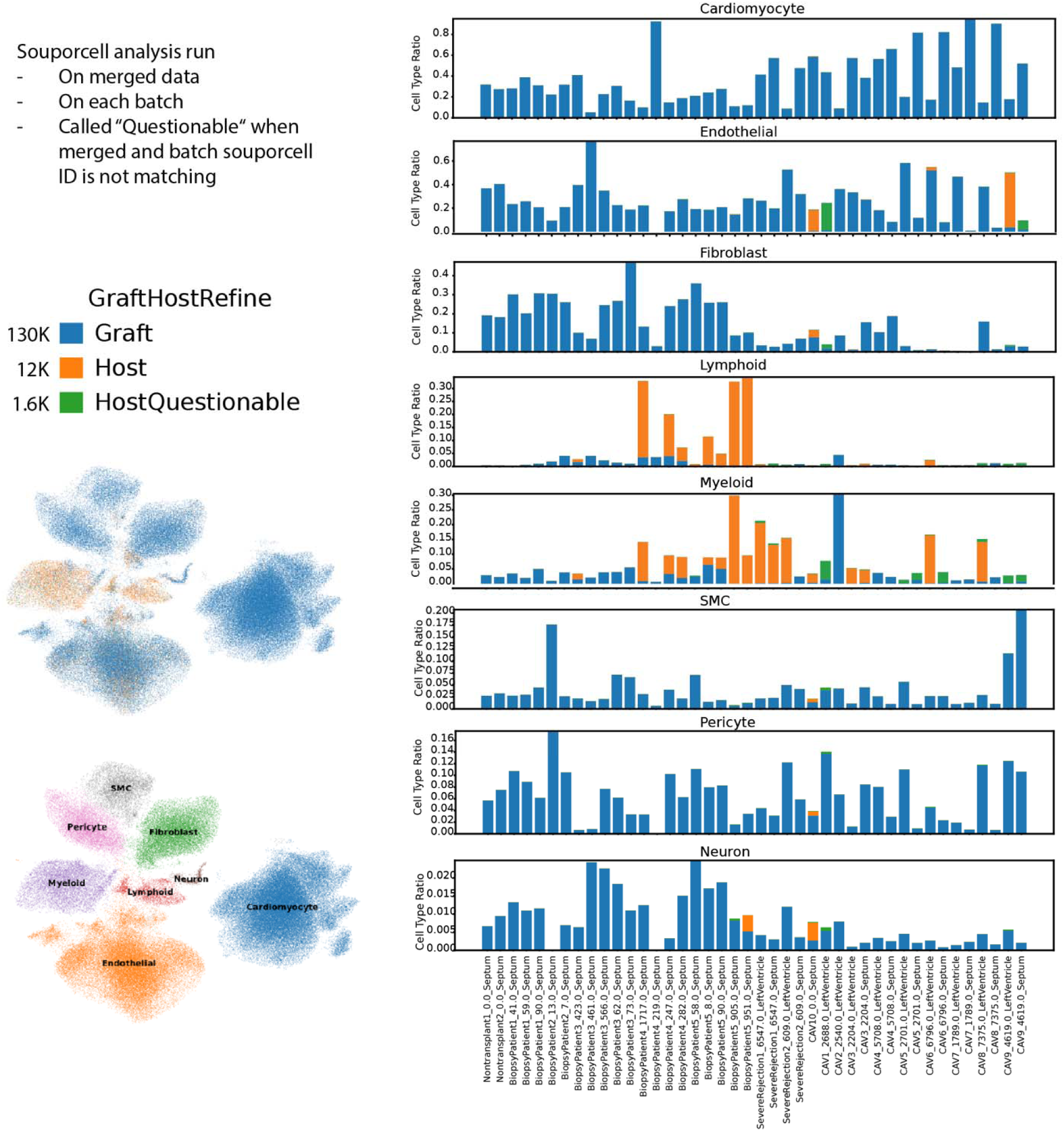
Analysis of graft versus host status by SouporCell de novo SNP analysis. Analysis of graft versus host status in single cell data by Souporcell. Analysis was run on each batch individually, and to provide more power and to extend one ID between batches, on all batches joined together. Questionable calls are those where call by batch and across all batches did not align. Bar plots show fraction of each within each population. Each sample is labeled by donor, day, and location.

**SI Figure 3.**
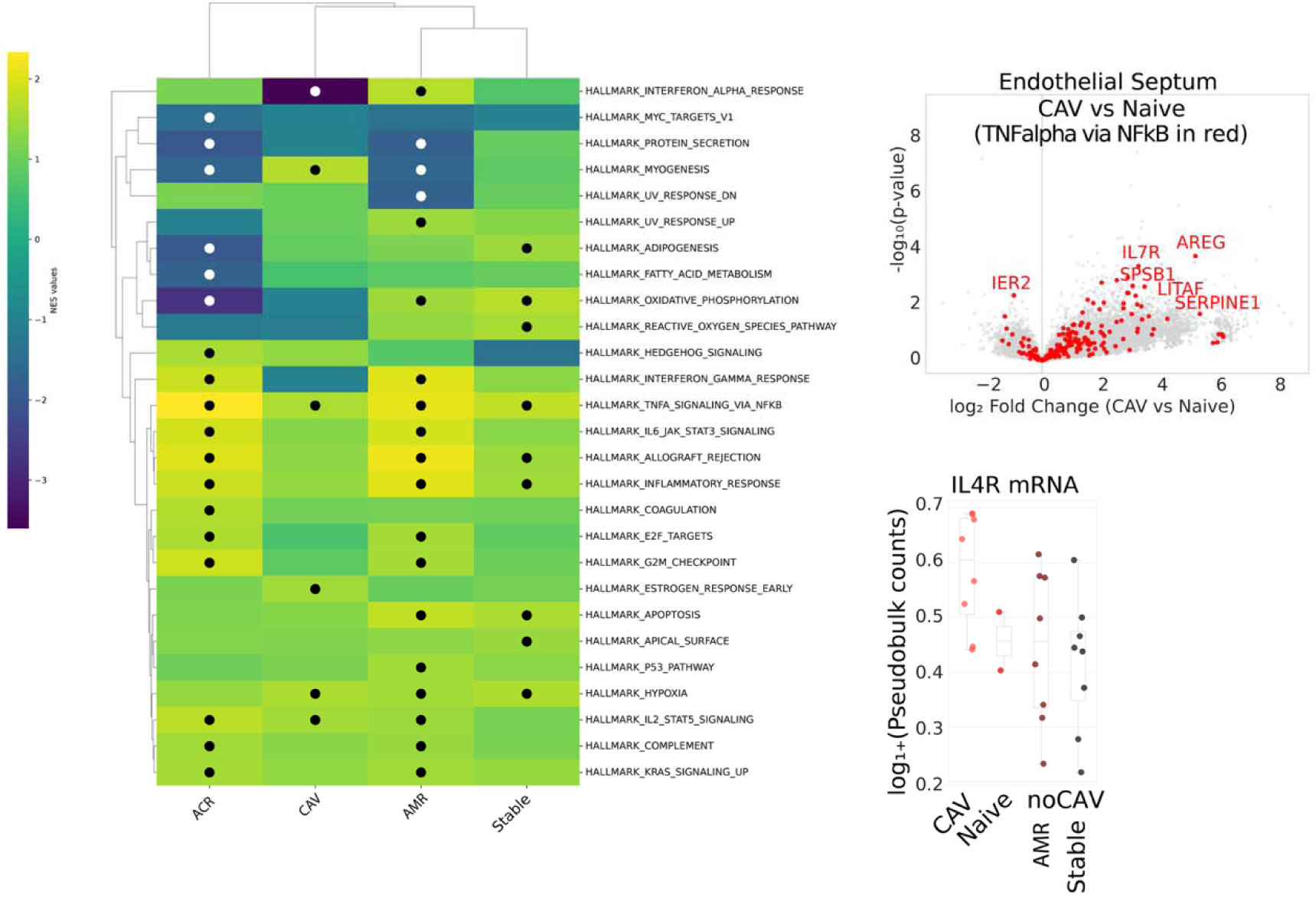
Gene set enrichment in endothelium, comparing samples to naïve heart endothelium. DESeq was used to examine differences between stable biopsies or unaffected heart tissues and CAV tissues for each cell population. Gene set enrichment was performed, using DESeq2 stat, to determine increased and decreased expression of gene sets. Circles = padj < 0.05. Volcano plot showing altered gene expression in the endothelial cell clusters (pseudobulk) in septal samples from CAV-positive explant versus naïve, with overlay of genes associated with the specific TNF-alpha mSigDB Hallmark pathway. Targeted analysis of IL4R showing differential expression, in quartile normalized read counts in each endothelial sample. CAV-negative biopsies were further separated into samples with mild rejection (Rejecting) or without signs of antibody or cellular rejection (Stable).

**SI Figure 4.**
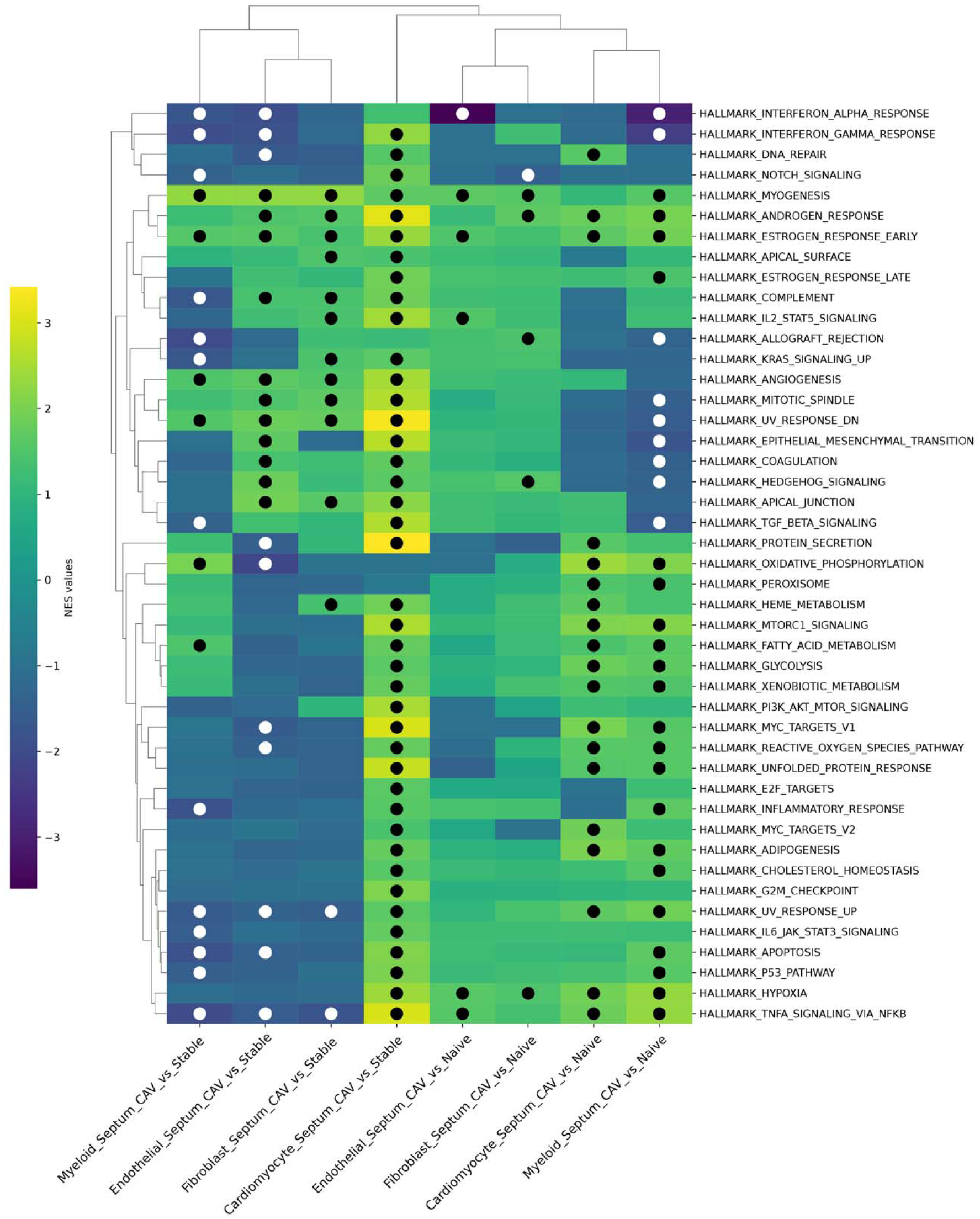
Gene set enrichment following pseudobulk analysis of all cells by major cell type. DESeq was used to examine differences between stable biopsies or unaffected heart tissues and CAV tissues for each cell population. Gene set enrichment was performed, using DESeq2 stat, to determine increased and decreased expression of gene sets. Circles = padj < 0.05.

**SI Figure 5.**
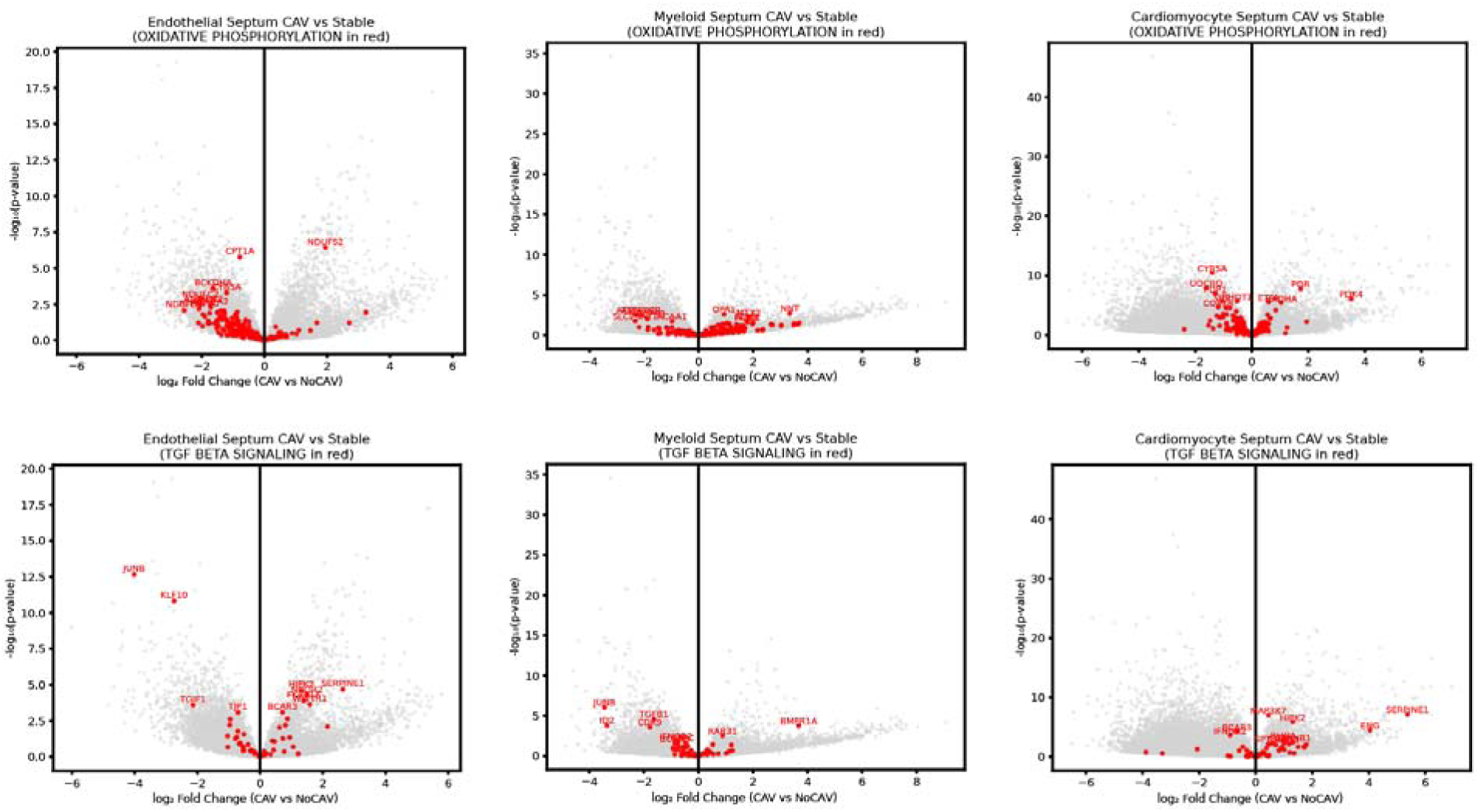
Volcano plots showing alterations in indicated gene sets. DEseq2 was used to determine altered expression patterns between stable biopsies the CAV tissues for each cell population. Red indicate the location of genes in the indicated pathways on these plots.

**SI Figure 6.**
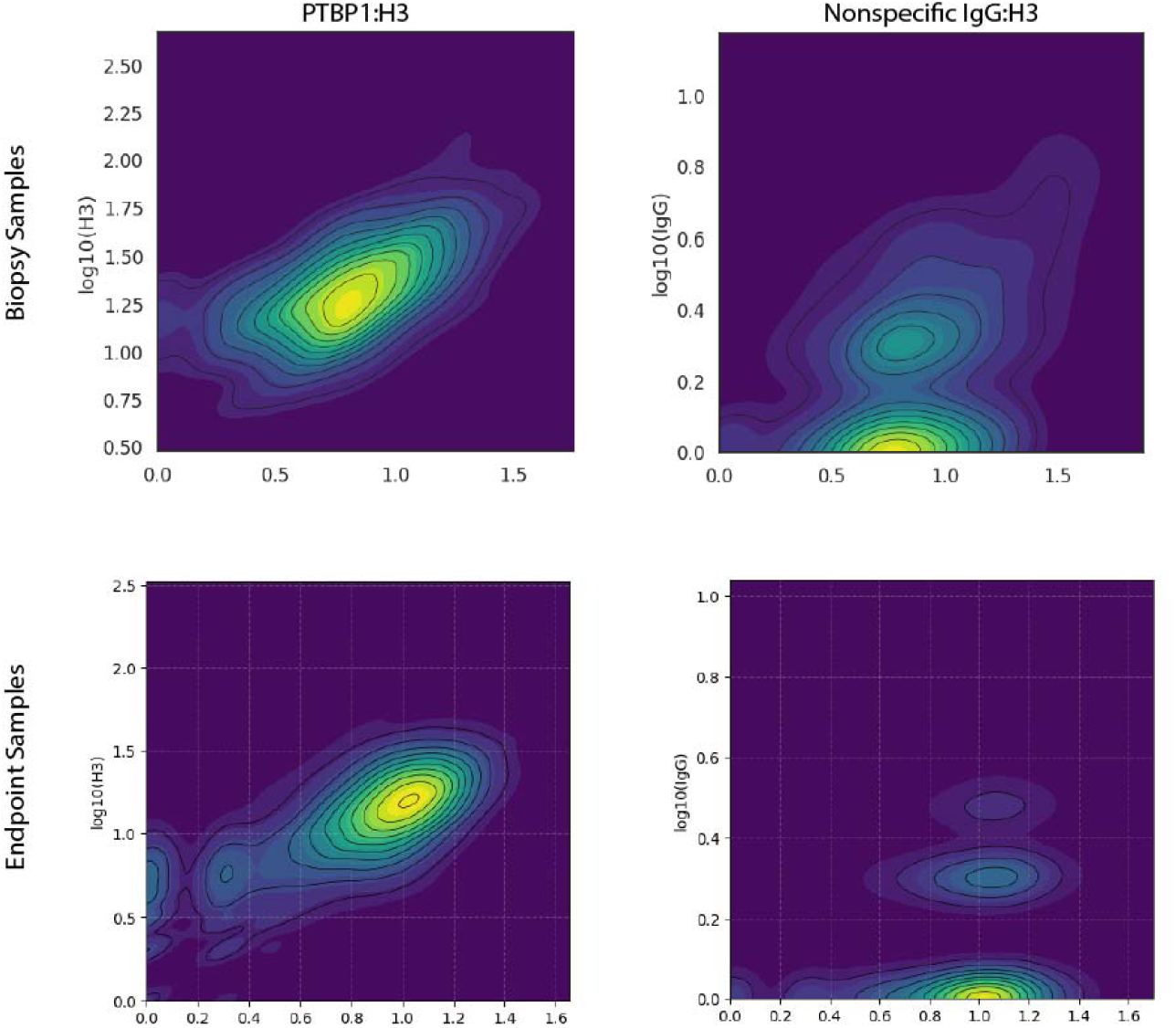
inCITE-seq protein counts. Raw protein counts per nuclei are show for each antibody. Percentages indicate the fraction of cells with 0 counts for the antibody. X-Y plots show the correlation between PTBP1 and histone H3 or IgG and histone H3.

**SI Figure 7.**
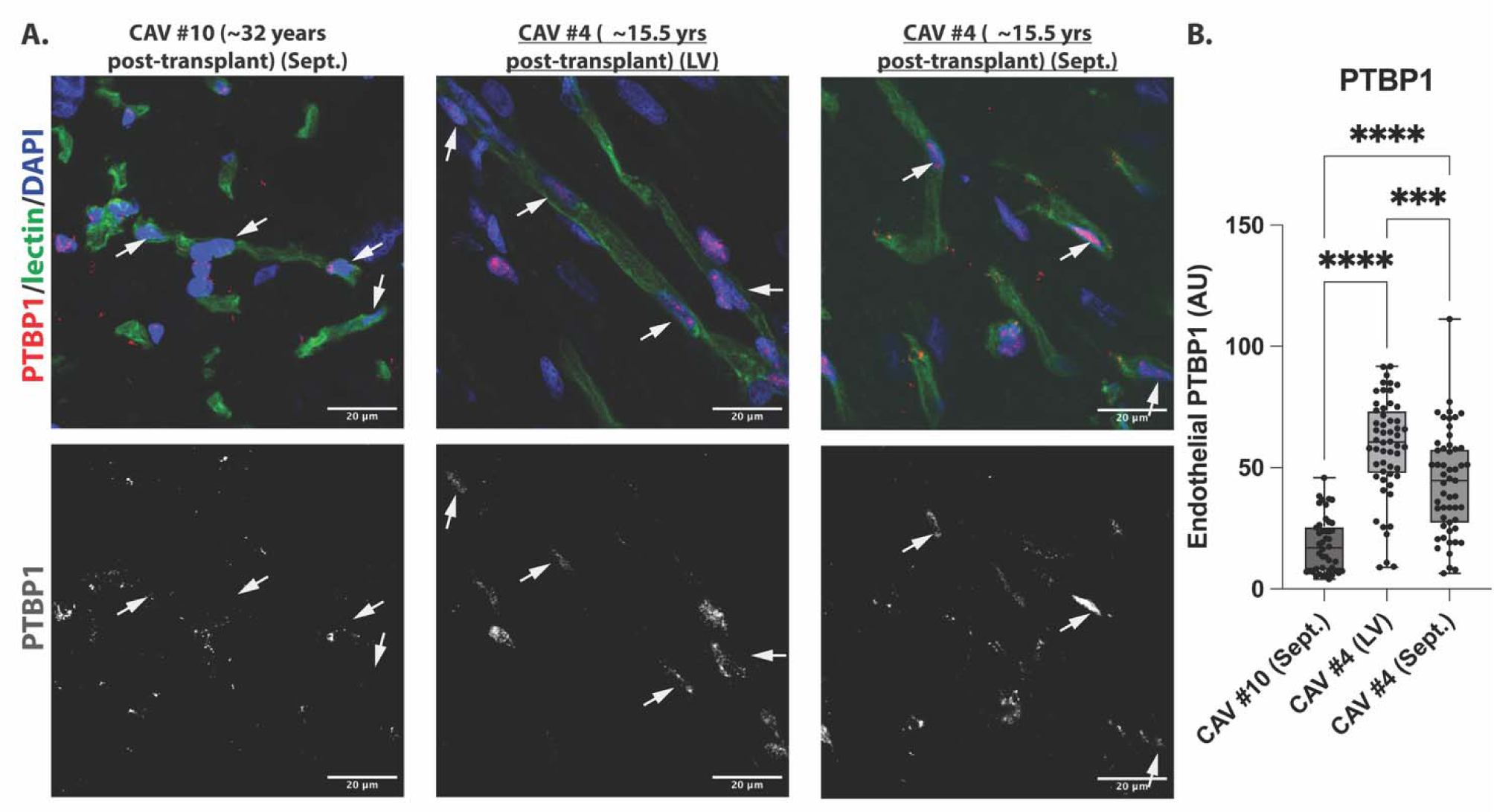
PTBP1 staining of long-lasting heart graft. (A) Immunofluorescence staining, showing PTBP1 levels in DAPI+ nuclei associated with UAE-1 lectin-stained vasculature (top panel). CAV tissues are from long-lasting CAV#10 sample (32 yrs post-transplant) or CAV#4 sample (15.5 years post-transplant) and either left ventricular (LV) or septal (Sept.). Bottom panel: grayscale PTBP1 staining alone (B) Quantification of endothelial PTBP1 levels across samples. Dots represent single cells from three random high-powered fields per sample. Statistical significance was assessed by one-way ANOVA with Tukey’s multiple comparisons test.

**SI Figure 8.**
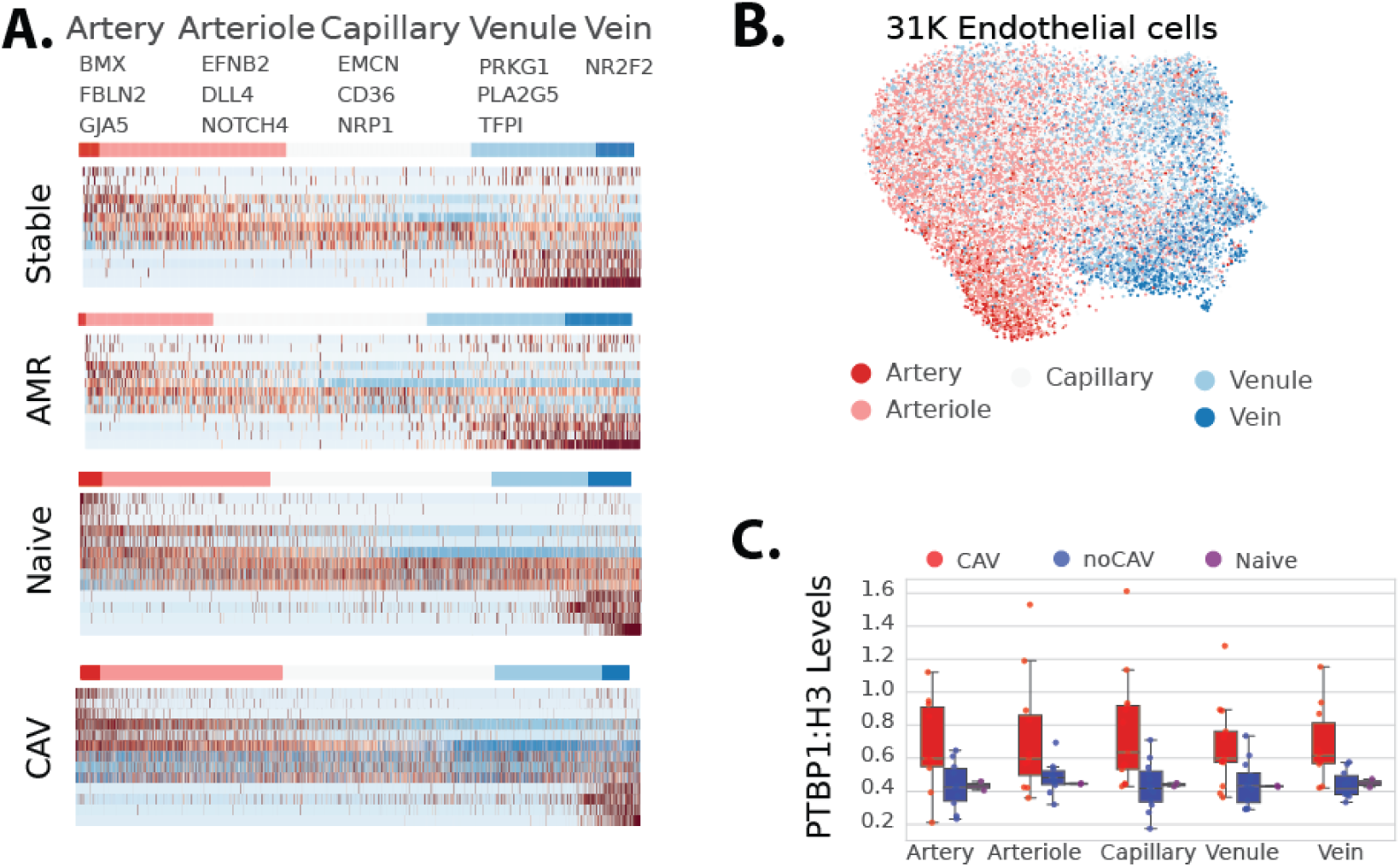
inCITE-seq PTBP1 counts across the AV-hierarchy. (A) Heat map aligning endothelial cells along arteriovenous heriarchy, based on scoring of arterial and venous genes (above). (B) Categorization of cells in capillary cluster. (C) PTBP1 counts, normalized to H3, from inCITE-seq data in cells within each region of the arteriovenous hierarchy.

**SI Figure 9.**
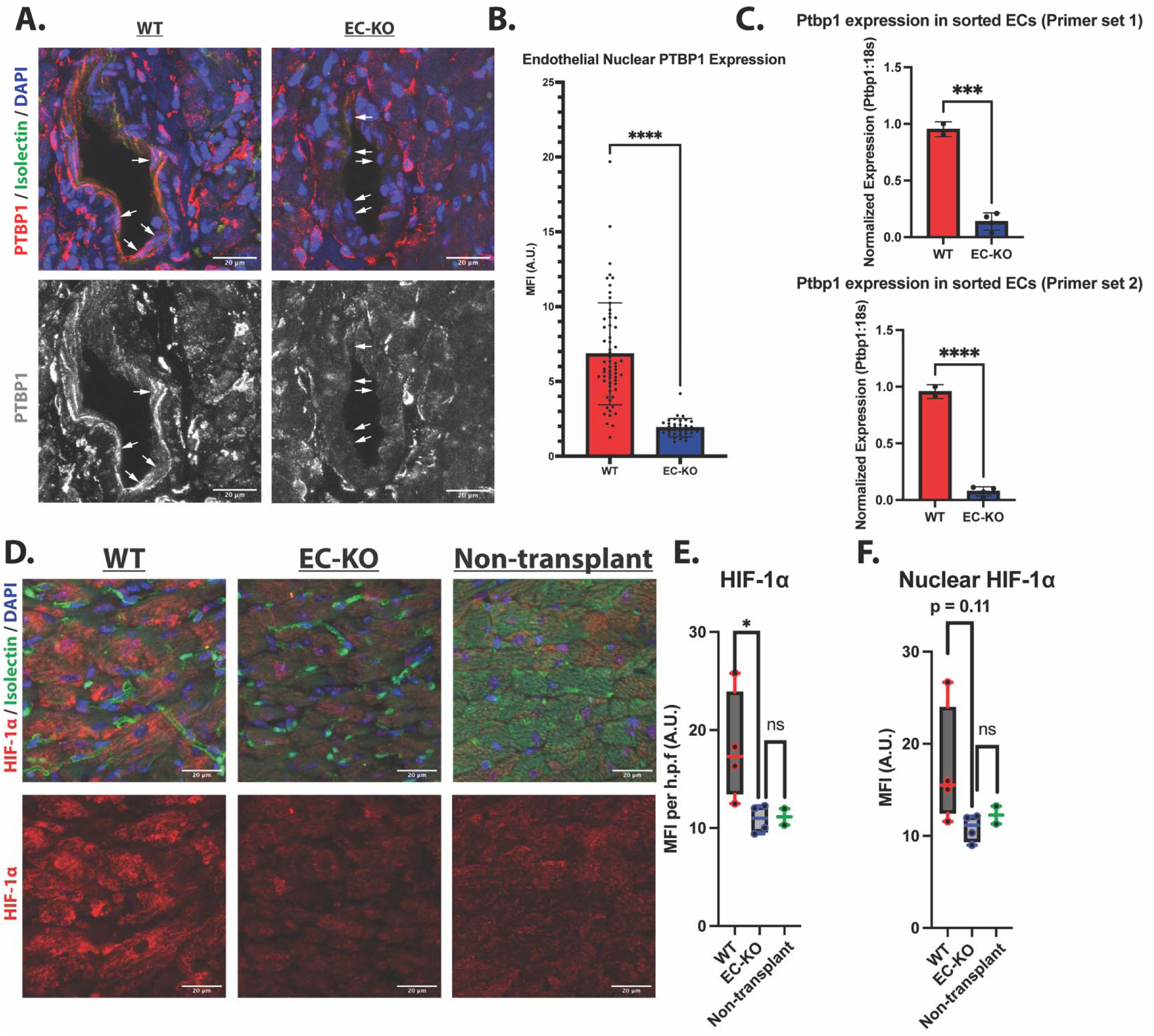
PTBP1 and HIF-1α staining of heart grafts. (A) Representative images of WT and EC-KO transplanted hearts stained for PTBP1 (red), isolectin B4 (green), and DAPI (blue). Bottom panels show grayscale single-channel PTBP1 images. (B) Quantification of endothelial nuclear PTBP1 expression in WT (n = 4) and EC-KO (n = 4) grafts. Each dot represents an individual endothelial cell quantified from three images per sample (>20 endothelial cells per sample). Data are shown as mean ± SD; significance was assessed using Mann-Whitney test (****p<0.0001). (C) Quantitative PCR analysis of Ptbp1 expression in CD31+ICAM2+ sorted endothelial cells isolated from WT (n = 2) and EC-KO (n = 4) grafts post-transplant, using two independent primer sets. Statistical significance was assessed by Student’s t test (D, E,F) Representative images (D) and quantification of overall mean fluorescence intensity (E) and nuclear mean fluorescence intensity (F) of HIF-1α in WT (n = 4) and EC-KO (n = 4) grafts and in non-transplanted control hearts (n = 2). Data are presented as median, min, and max. Statistical significance was determined using Mann-Whitney test (*p < 0.05).

**SI Figure 10.**
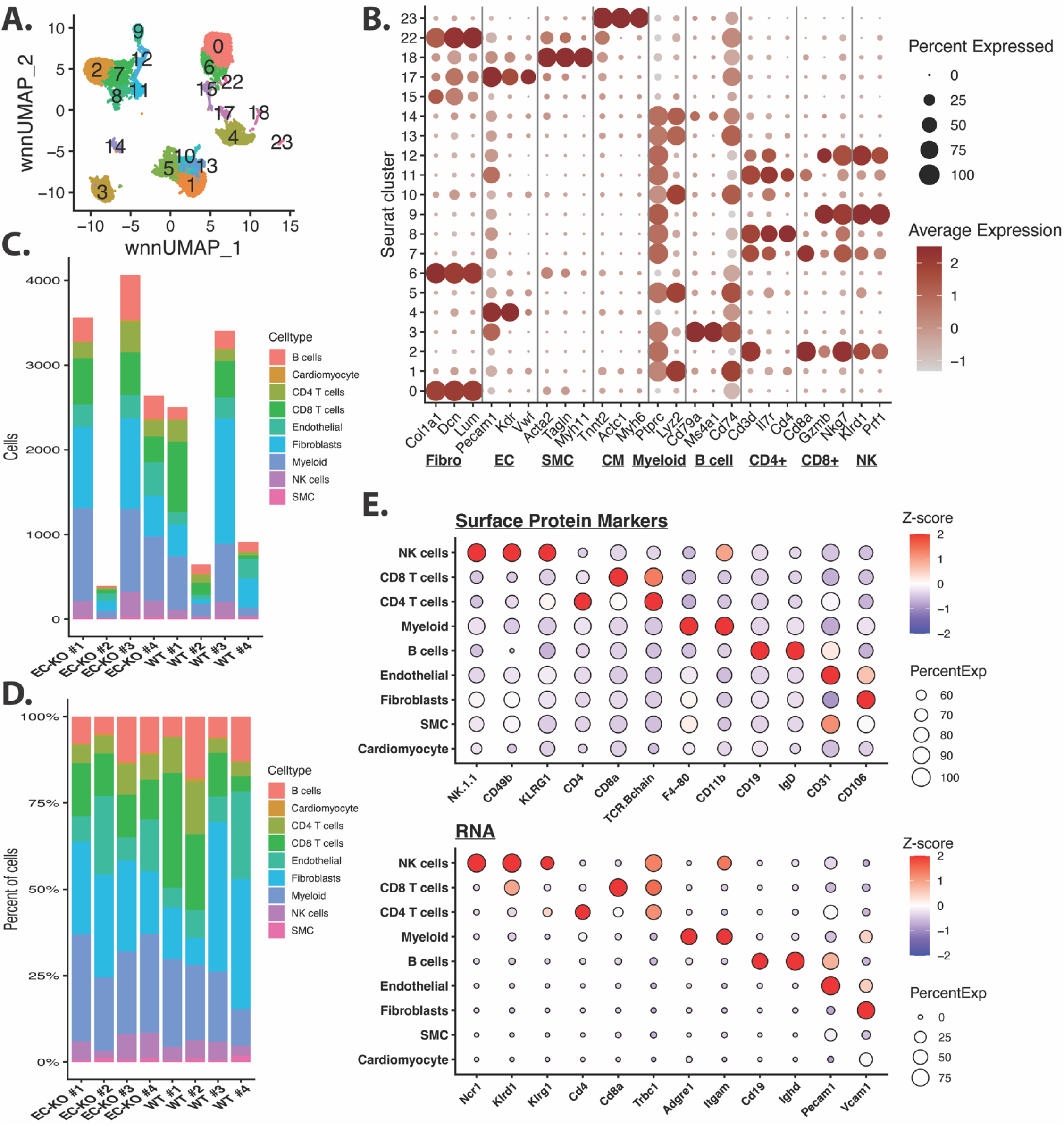
Cardiac cell composition of WT and EC-KO grafts by CITE-seq. (A) Weighted nearest neighbor (WNN) RNA–protein integrated UMAP depicting cardiac cell clusters identified by CITE-seq. (B) Dot plot showing canonical RNA markers used to annotate cardiac cell clusters. (C) Number of cells per sample, stratified by cell type. (D) Proportion of each cardiac cell type within individual samples. (E) Dot plot displaying RNA and surface protein expression of canonical markers across major cardiac cell types.

**SI Figure 11.**
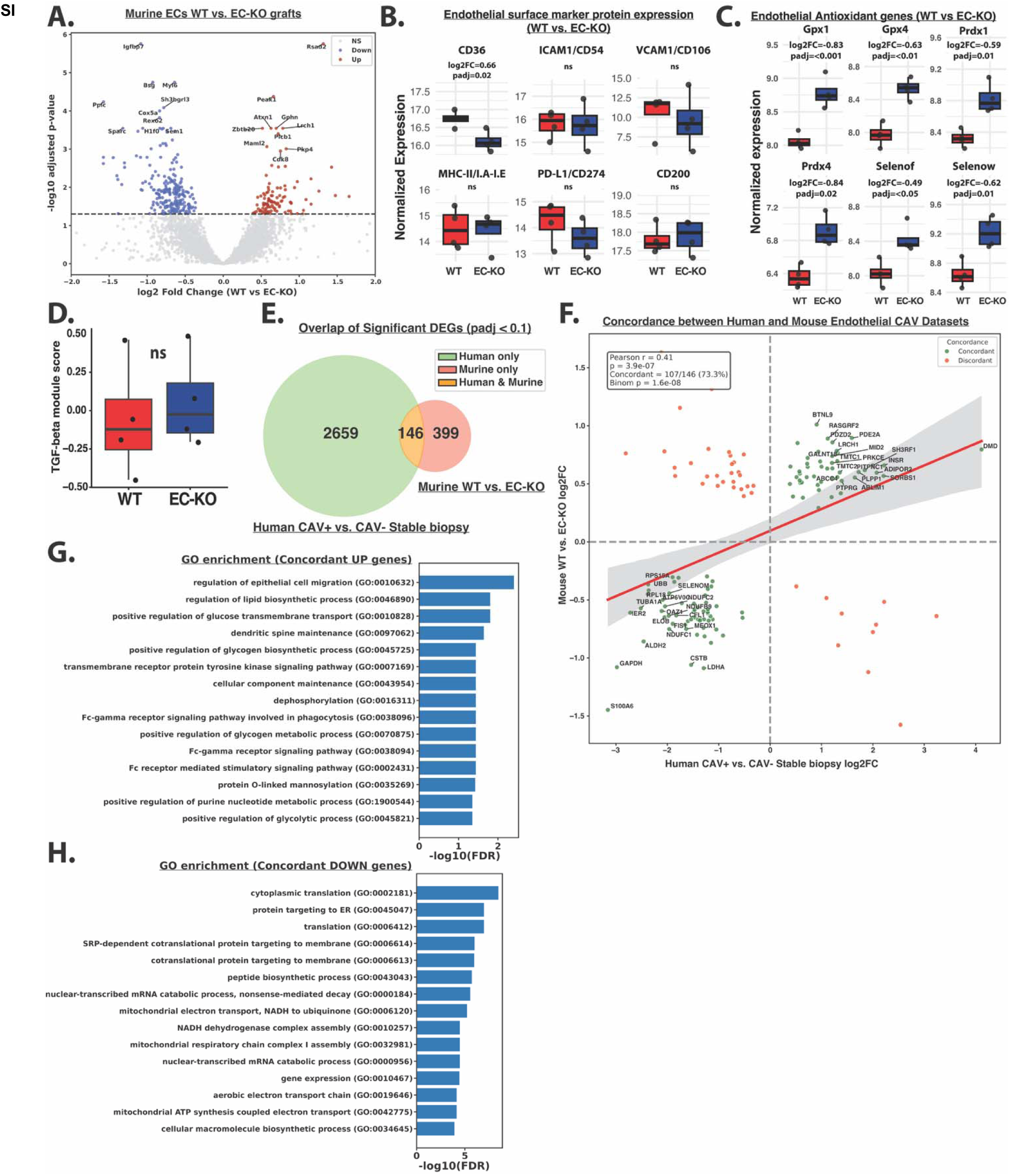
Endothelial cell transcriptional analysis. (A) Volcano plot of pseudobulk differentially expressed genes (DEGs) comparing WT and EC-KO endothelial cells. Significantly upregulated genes are shown in red and downregulated in blue. (B) Pseudobulk expression of select surface protein markers in endothelial cells from WT and EC-KO grafts. Bar plots represent variance-stabilizing transformation (VST)-normalized expression, log2FC and adjusted p-values were calculated using DESeq2. (C) Pseudobulk expression of select antioxidant genes in endothelial cells from WT and EC-KO grafts. Bar plots represent variance-stabilizing transformation (VST)-normalized expression, log2FC and adjusted p-values were calculated using DESeq2. (D) Pseudobulk module score of Hallmark TGF-β response gene set between WT and EC-KO grafts. Statistical significance was assessed using the Mann-Whitney U test. (E) Overlap of significantly differentially expressed genes in human (CAV+ vs. CAV- stable biopsy) and murine (WT vs. EC-KO) endothelial datasets. (F) Scatter plot of the 146 shared differentially expressed genes shared between the human and mouse EC datasets. Concordant genes are changing in the same direction in both datasets. Significance of concordance was tested using a binomial test (null hypothesis= 50% concordance), and Pearson correlation was used to assess correlation of gene expression changes across species. (G) Gene ontology (GO) enrichment analysis of concordant upregulated genes between human and mouse datasets. (H) GO enrichment analysis of concordant downregulated genes between human and mouse datasets.

**SI Figure 12.**
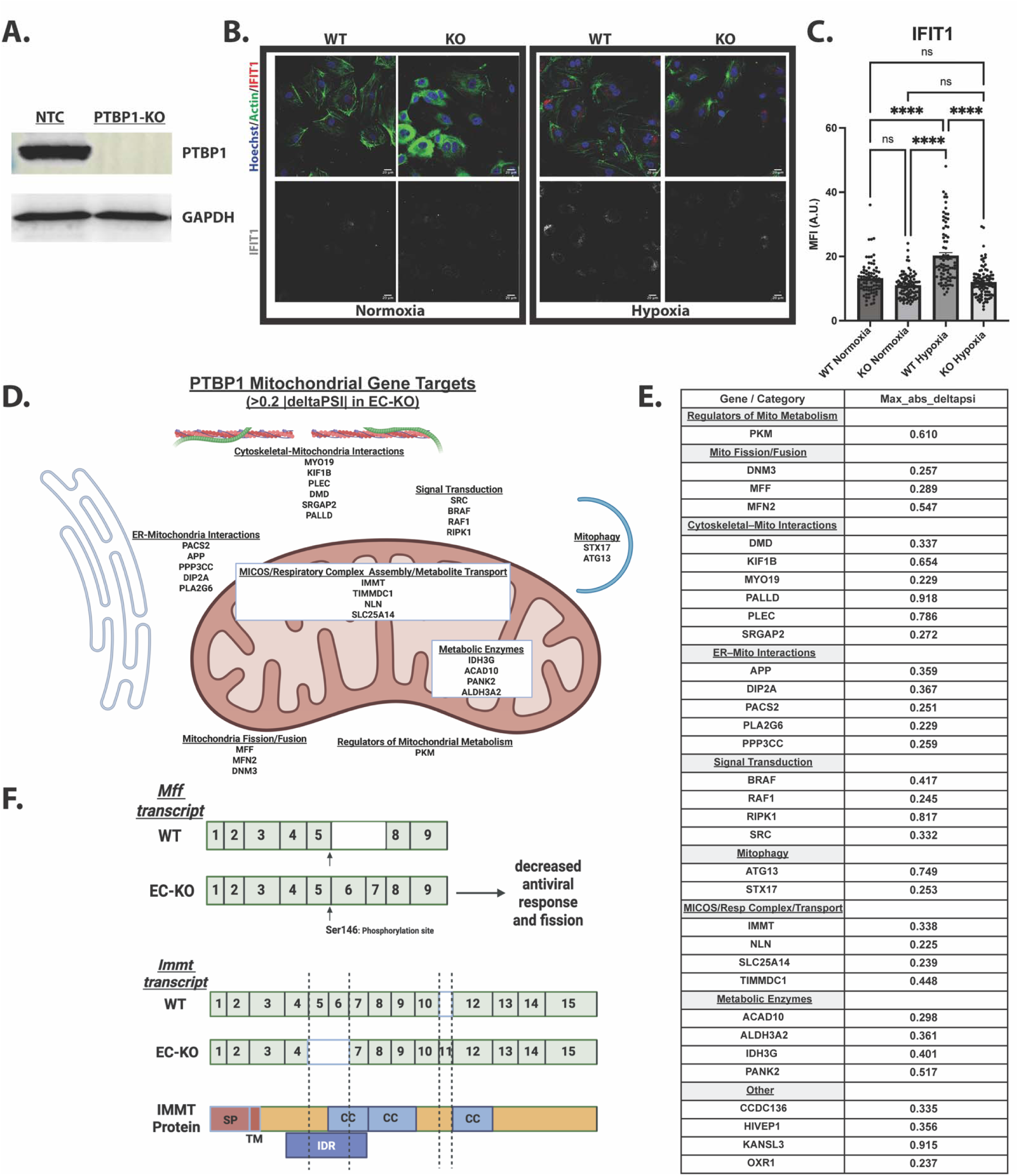
PTBP1 mediates splicing of endothelial mitochondrial transcripts. (A) Immunoblot of PTBP1 and GAPDH in non-targeting control (NTC) and PTBP1-knockout (KO) pooled HUVECs (B) Representative immunofluorescence images of WT and PTBP1 KO HUVECs cultured +/- hypoxia for 48 hrs and stained with IFIT1, phalloidin (actin), and Hoechst (upper panel). IFIT1 staining alone (grayscale) (bottom panel). (C) Quantification of IFIT1 fluorescence intensity. Each dot represents an individual cell from two biological replicates. Statistical significance was determined by one-way ANOVA with Tukey’s multiple comparisons test. (D) Schematic overview of mitochondrial genes exhibiting significant alternative splicing (|deltaPSI| >0.2) in PTBP1 EC-KO murine endothelial cells. (E) Table of PTBP1-mediated spliced mitochondrial genes in isolated murine arterial endothelial cells shown as ΔPSI values. Data were reanalyzed from (Hensel *et al*., 2022). (F) Gene models showing alternative splicing changes in Mff and Immt gene upon EC-KO. Alignment of Uniprot protein domains with exons was performed using UCSC Genome Browser. SP: signal peptide, TM: transmembrane domain, IDR: intrinsically disordered domain, CC: coiled-coil domain. Schematics in (D) and (F) were made using BioRender.

**SI Figure 13.**
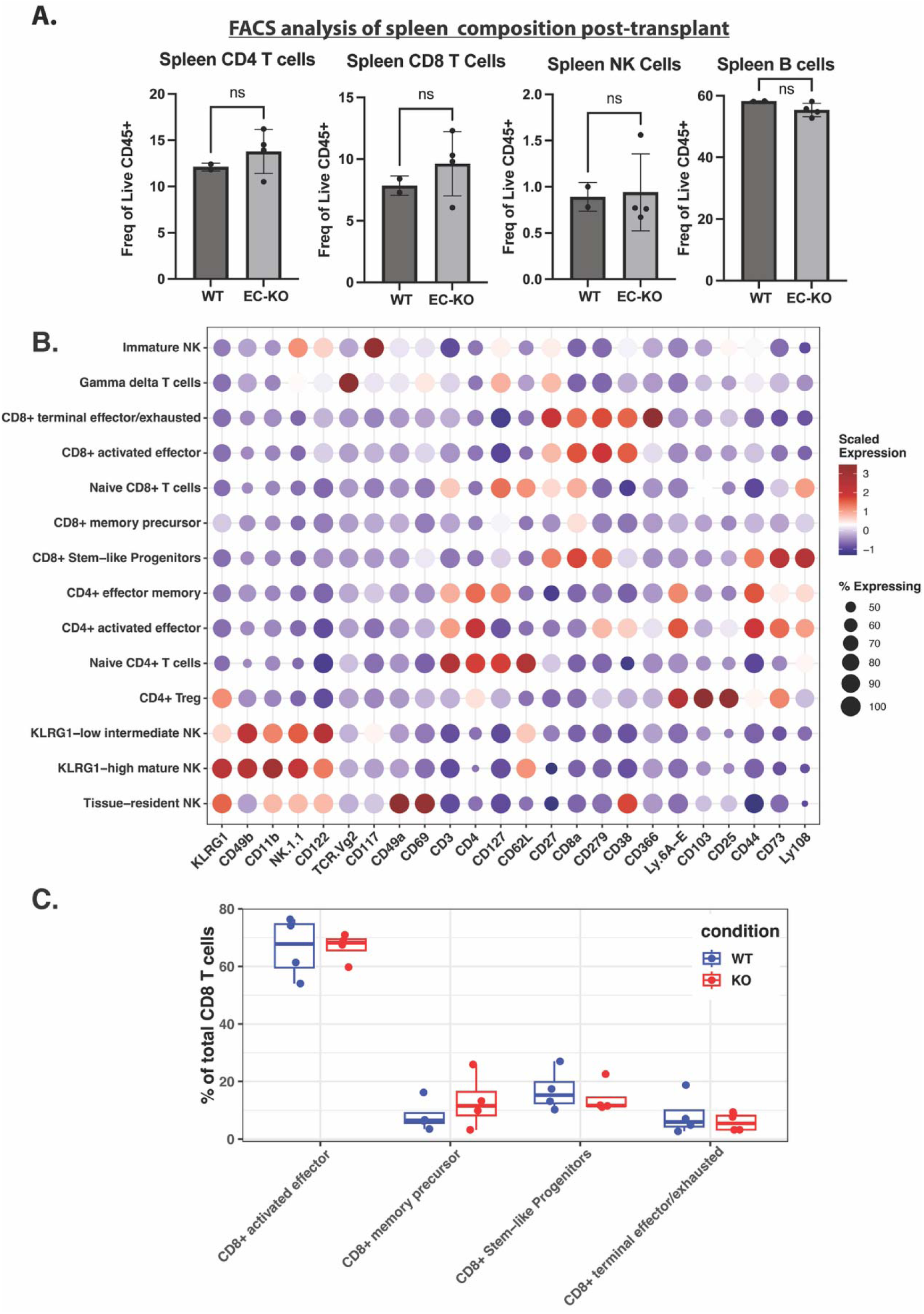
Spleen composition post-transplant and identification of lymphocyte subclusters in WT and EC-KO grafts using CITE-seq. (A) Flow cytometric quantification of CD4+ T cells, CD8+ T cells, NK1.1+ NK cells, and CD19+ B cells in spleens following transplantation (WT, n = 2; EC-KO, n = 4). Data are shown as mean ± SD; significance was assessed using an unpaired two-tailed Student’s t test.( B) Dot plot of surface protein markers used to distinguish T and NK subtypes. Immature NK (NK.1.1+, CD122+, CD117+), Tissue-resident NK (NK1.1+, KLRG1+, CD49a+, CD69+, CD49b-), KLRG1-low intermediate NK (NK1.1+, KLRG1 low, CD11b+, CD49b+, CD122+), KLRG1-high mature NK (NK1.1+, KLRG1 high, CD11b+, CD49b+, CD62L high), Naïve CD4+ T cells (CD3+, CD4+, CD127+, CD62L+), CD4+ effector memory (CD3+, CD4+, CD44+), CD4+ activated effector (CD3+, CD4+, CD44+, Ly6A-E high, CD279+, CD38+, CD73+), CD4+ Treg (CD4+, CD103+, CD25+, KLRG1+), Naïve CD8+ T cells (CD3+, CD8a+, CD127+, CD62L+), CD8+ memory precursor (CD8a+, CD127 low, CD62L-), CD8+ stem-like progenitor T cell (CD8a+, CD27+, CD279+, CD44+, CD73+,Ly108+), CD8+ activated effector T cells (CD8a+, CD27+ CD38+, CD279+), CD8+ terminal effector/exhausted T cells ( CD8a+, CD27+ CD38+, CD279+, CD366+), Gamma-delta (TCR-VG+). (C). Proportion of CD8+ T cell subsets of total T cells per sample and condition (WT, n = 4, EC-KO, n =4).

**SI Figure 14.**
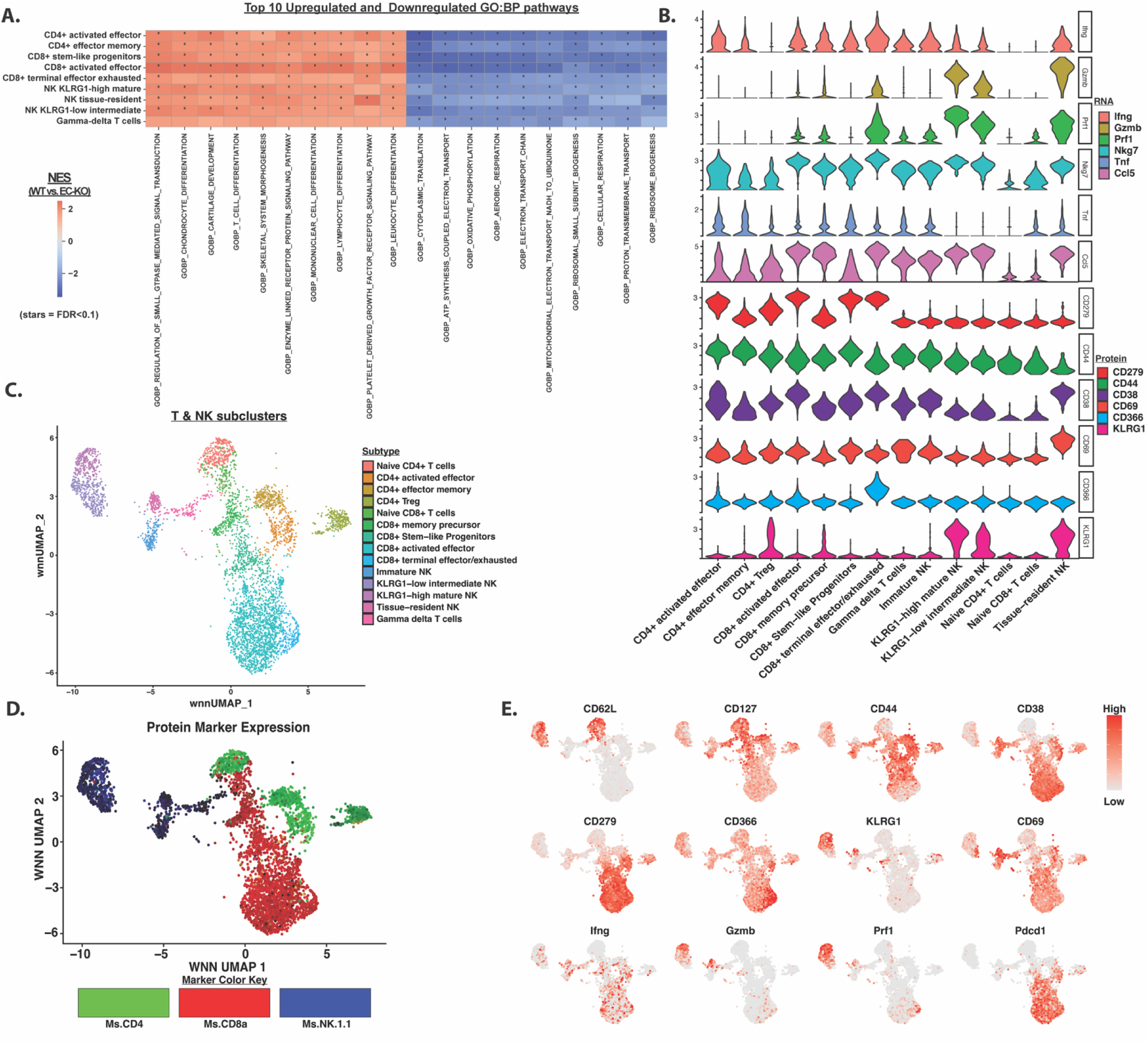
Differential immune signaling in T and NK subsets between WT and EC-KO grafts. (A) Heatmap showing the top 10 upregulated and downregulated gene ontology (GO) pathways in T and NK subsets in WT vs. EC-KO grafts. Normalized enrichment score (NES), stars denote pathways with FDR q value <0.1. (B) Violin plots of RNA and protein marker expression across T and NK cell subtypes. (C) UMAP of T and NK subtypes. (D) Feature plot of CD4, CD8a, and NK1.1 protein expression. (E) Feature plots of protein and RNA markers.

## SI Tables

For processing details see : https://github.com/pamurphyUCONN/2026_Pathoulas

**SI Table 1.**
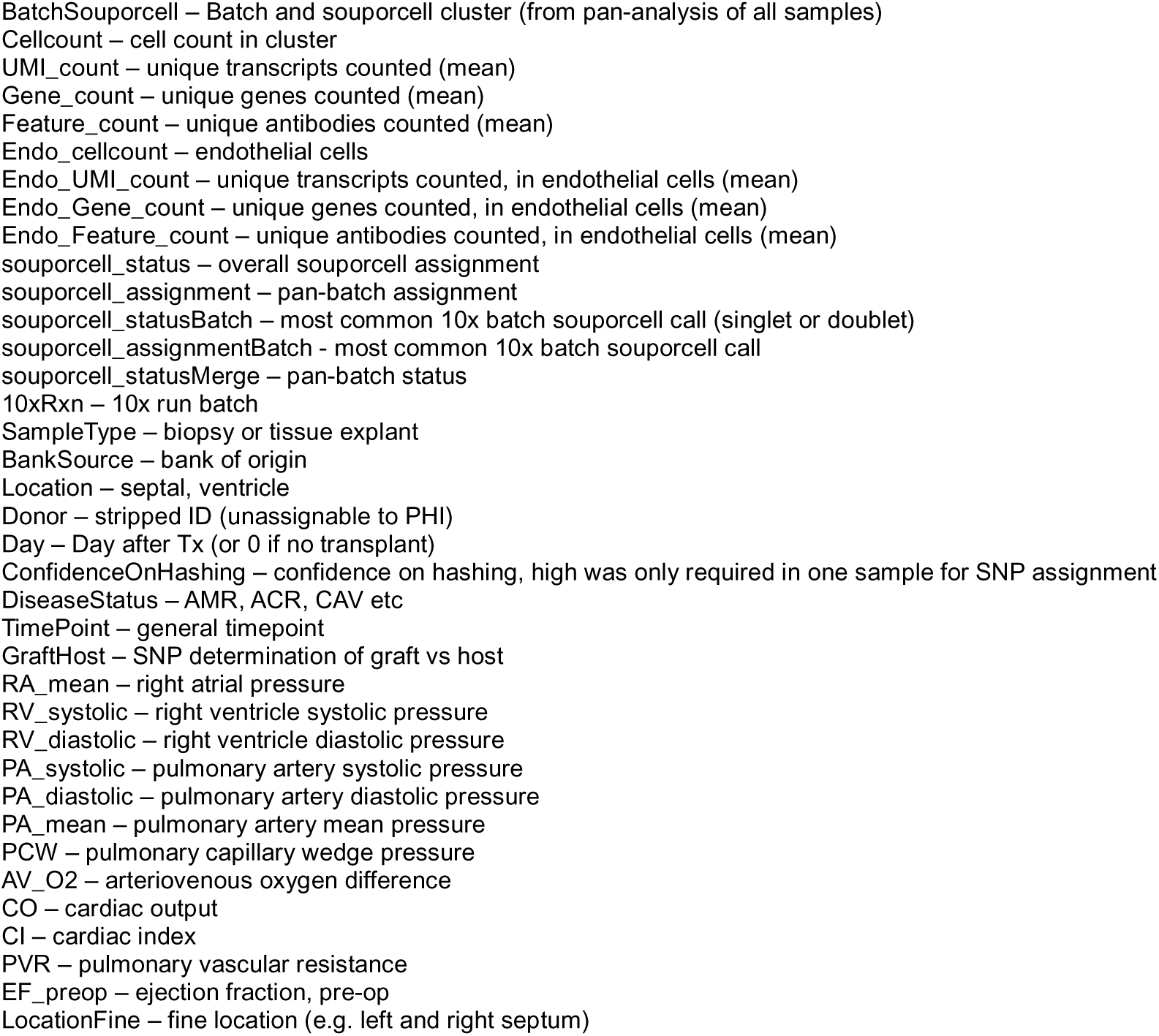
Donor Tissues.

SI Table 2. Pseudobulk DESeq2 analysis by major cell type and disease state, human heart tissues.

SI Table 3A-B. Pseudobulk GSEA analysis by major cell type and disease state, human heart tissues.

SI Table 4A-D. Pseudobulk DESeq2 analysis by nuclear PTBP1 levels, human heart tissues.

SI Table 5A-D. Pseudobulk GSEA analysis by nuclear PTBP1 levels, human heart tissues.

SI Table 6. CITE-seq surface protein markers

SI Table 7. Pseudobulk mRNA and surface protein DESeq2 analysis by major cell type comparing genotypes, murine heart tissues.

SI Table 8. Pseudobulk GSEA analysis by major cell type comparing genotypes, murine heart tissues.

SI Table 9. Concordant genes between human (CAV+ vs. CAV- stable biopsy) and murine (WT vs. EC-KO) endothelial datasets.

SI Table 10. GSEA analysis of concordant genes between human (CAV+ vs. CAV- stable biopsy) and murine (WT vs. EC-KO) endothelial datasets.

SI Table 11. Pseudobulk mRNA DESeq2 analysis in T and NK subtypes comparing genotypes, murine heart tissues.

SI Table 12. Pseudobulk GSEA analysis in T and NK subtypes comparing genotypes, murine heart tissues.

SI Table 13. Pseudobulk protein analysis in T and NK subtypes comparing genotypes, murine heart tissues.

SI Table 14. Key reagents.

## References

1. Pober JS, Chih S, Kobashigawa J, Madsen JC, Tellides G. Cardiac allograft vasculopathy: current review and future research directions. Cardiovasc Res. 2021 Nov 22;117(13):2624–2638. PMCID: PMC8783389

2. Rahmani M, Cruz RP, Granville DJ, McManus BM. Allograft Vasculopathy Versus Atherosclerosis. Circ Res. American Heart Association; 2006 Oct 13;99(8):801–815.

3. Chen PY, Qin L, Baeyens N, Li G, Afolabi T, Budatha M, Tellides G, Schwartz MA, Simons M. Endothelial-to-mesenchymal transition drives atherosclerosis progression. J Clin Invest. 2015 Oct 26;125(12):4514–4528. PMCID: PMC4665771

4. Chen PY, Qin L, Li G, Wang Z, Dahlman JE, Malagon-Lopez J, Gujja S, Cilfone NA, Kauffman KJ, Sun L, Sun H, Zhang X, Aryal B, Canfran-Duque A, Liu R, Kusters P, Sehgal A, Jiao Y, Anderson DG, Gulcher J, Fernandez-Hernando C, Lutgens E, Schwartz MA, Pober JS, Chittenden TW, Tellides G, Simons M. Endothelial TGF-β signalling drives vascular inflammation and atherosclerosis. Nat Metab. 2019 Sept;1(9):912–926. PMCID: PMC6767930

5. Zou H, Ming B, Li J, Xiao Y, Lai L, Gao M, Xu Y, Tan Z, Gong F, Zheng F. Extracellular HMGB1 Contributes to the Chronic Cardiac Allograft Vasculopathy/Fibrosis by Modulating TGF-β1 Signaling. Front Immunol [Internet]. 2021;Volume 12-2021. Available from: https://www.frontiersin.org/journals/immunology/articles/10.3389/fimmu.2021.641973

6. Sun X, Lu Q, Yegambaram M, Kumar S, Qu N, Srivastava A, Wang T, Fineman JR, Black SM. TGF-β1 attenuates mitochondrial bioenergetics in pulmonary arterial endothelial cells via the disruption of carnitine homeostasis. Redox Biol. 2020 Sept 1;36:101593.

7. Pardali E, Sanchez-Duffhues G, Gomez-Puerto MC, Ten Dijke P. TGF-β-Induced Endothelial-Mesenchymal Transition in Fibrotic Diseases. Int J Mol Sci. 2017;18(10):2157.

8. Hong SG, Shin J, Choi SY, Powers JC, Meister BM, Sayoc J, Son JS, Tierney R, Recchia FA, Brown MD, Yang X, Park JY. Flow pattern-dependent mitochondrial dynamics regulates the metabolic profile and inflammatory state of endothelial cells. JCI Insight. 2022 Sept 22;7(18):e159286. PMCID: PMC9514384

9. Forrester SJ, Preston KJ, Cooper HA, Boyer MJ, Escoto KM, Poltronetti AJ, Elliott KJ, Kuroda R, Miyao M, Sesaki H, Akiyama T, Kimura Y, Rizzo V, Scalia R, Eguchi S. Mitochondrial Fission Mediates Endothelial Inflammation. Hypertension. American Heart Association; 2020 July 1;76(1):267–276.

10. Tran DT, Tu Z, Alawieh A, Mulligan J, Esckilsen S, Quinn K, Sundararaj K, Wallace C, Finnegan R, Allen P, Mehrotra S, Atkinson C, Nadig SN. Modulating donor mitochondrial fusion/fission delivers immunoprotective effects in cardiac transplantation. Am J Transplant. John Wiley & Sons, Ltd; 2022 Feb 1;22(2):386–401.

11. Hensel JA, Nicholas SAE, Kimble AL, Nagpal AS, Omar OMF, Tyburski JD, Jellison ER, Ménoret A, Ozawa M, Rodriguez-Oquendo A, Vella AT, Murphy PA. Splice factor polypyrimidine tract-binding protein 1 (Ptbp1) primes endothelial inflammation in atherogenic disturbed flow conditions. Proc Natl Acad Sci U S A. United States; 2022 July 26;119(30):e2122227119. PMCID: PMC9335344

12. Caruso P, Dunmore BJ, Schlosser K, Schoors S, Dos Santos C, Perez-Iratxeta C, Lavoie JR, Zhang H, Long L, Flockton AR, Frid MG, Upton PD, D’Alessandro A, Hadinnapola C, Kiskin FN, Taha M, Hurst LA, Ormiston ML, Hata A, Stenmark KR, Carmeliet P, Stewart DJ, Morrell NW. Identification of MicroRNA-124 as a Major Regulator of Enhanced Endothelial Cell Glycolysis in Pulmonary Arterial Hypertension via PTBP1 (Polypyrimidine Tract Binding Protein) and Pyruvate Kinase M2. Circulation. American Heart Association; 2017 Dec 19;136(25):2451–2467.

13. Liu H, Duan R, He X, Qi J, Xing T, Wu Y, Zhou L, Wang L, Shao Y, Zhang F, Zhou H, Gu X, Lin B, Liu Y, Wang Y, Liu Y, Li L, Liang D, Chen YH. Endothelial deletion of PTBP1 disrupts ventricular chamber development. Nat Commun. 2023 Mar 31;14(1):1796. PMCID: PMC10066379

14. Caruso P, Dunmore BJ, Schlosser K, Schoors S, Dos Santos C, Perez-Iratxeta C, Lavoie JR, Zhang H, Long L, Flockton AR, Frid MG, Upton PD, D’Alessandro A, Hadinnapola C, Kiskin FN, Taha M, Hurst LA, Ormiston ML, Hata A, Stenmark KR, Carmeliet P, Stewart DJ, Morrell NW. Identification of MicroRNA-124 as a Major Regulator of Enhanced Endothelial Cell Glycolysis in Pulmonary Arterial Hypertension via PTBP1 (Polypyrimidine Tract Binding Protein) and Pyruvate Kinase M2. Circulation. 2017 Dec 19;136(25):2451–2467. PMCID: PMC5736425

15. Omar OMF, Kimble AL, Cheemala A, Tyburski JD, Pandey S, Wu Q, Reese B, Jellison ER, Hao B, Li Y, Yan R, Murphy PA. Endothelial TDP-43 depletion disrupts core blood–brain barrier pathways in neurodegeneration. Nat Neurosci. 2025 May 1;28(5):973–984.

16. Amancherla K, Qin J, Hulke ML, Pfeiffer RD, Agrawal V, Sheng Q, Xu Y, Schlendorf KH, Lindenfeld J, Shah RV, Freedman JE, Tucker NR, Moslehi J. Single-Nuclear RNA Sequencing of Endomyocardial Biopsies Identifies Persistence of Donor-Recipient Chimerism With Distinct Signatures in Severe Cardiac Allograft Vasculopathy. Circ Heart Fail. United States; 2023 Jan;16(1):e010119. PMCID: PMC9852032

17. Quaini Federico, Urbanek Konrad, Beltrami Antonio P., Finato Nicoletta, Beltrami Carlo A., Nadal-Ginard Bernardo, Kajstura Jan, Leri Annarosa, Anversa Piero. Chimerism of the Transplanted Heart. N Engl J Med. Massachusetts Medical Society; 346(1):5–15.

18. Kindel SJ, Law YM, Chin C, Burch M, Kirklin JK, Naftel DC, Pruitt E, Carboni MP, Arens A, Atz AM, Dreyer WJ, Mahle WT, Pahl E. Improved Detection of Cardiac Allograft Vasculopathy: A Multi-Institutional Analysis of Functional Parameters in Pediatric Heart Transplant Recipients. J Am Coll Cardiol. 2015 Aug 4;66(5):547–557.

19. Tona F, Osto E, Famoso G, Previato M, Fedrigo M, Vecchiati A, Perazzolo Marra M, Tellatin S, Bellu R, Tarantini G, Feltrin G, Angelini A, Thiene G, Gerosa G, Iliceto S. Coronary Microvascular Dysfunction Correlates With the New Onset of Cardiac Allograft Vasculopathy in Heart Transplant Patients With Normal Coronary Angiography. Am J Transplant. 2015 May 1;15(5):1400–1406.

20. Chih S, Chong AY, Džavík V, So DY, Aleksova N, Wells GA, Bernick J, Overgaard CB, Stadnick E, Mielniczuk LM, Beanlands RSB, Ross HJ. Fibrotic Plaque and Microvascular Dysfunction Predict Early Cardiac Allograft Vasculopathy Progression After Heart Transplantation: The Early Post Transplant Cardiac Allograft Vasculopathy Study. Circ Heart Fail. American Heart Association; 2023 June 1;16(6):e010173.

21. Aziz T, Hasleton P, Hann AW, Yonan N, Deiraniya A, Hutchinson IV. Transforming growth factor β in relation to cardiac allograft vasculopathy after heart transplantation. J Thorac Cardiovasc Surg. 2000 Apr 1;119(4):700–708.

22. Aziz TM, Burgess MI, Haselton PS, Yonan NA, Hutchinson IV. Transforming growth factor β and diastolic left ventricular dysfunction after heart transplantation: echocardiographic and histologic evidence. J Heart Lung Transplant. 2003 June 1;22(6):663–673.

23. Ramzy D, Rao V, Brahm J, Miriuka S, Delgado D, Ross HJ. Cardiac allograft vasculopathy: a review. Can J Surg. 2005 Aug 1;48(4):319.

24. Redmond EM, Hamm K, Cullen JP, Hatch E, Cahill PA, Morrow D. Inhibition of Patched-1 Prevents Injury-Induced Neointimal Hyperplasia. Arterioscler Thromb Vasc Biol. American Heart Association; 2013 Aug 1;33(8):1960–1964.

25. Morrow D, Cullen JP, Liu W, Guha S, Sweeney C, Birney YA, Collins N, Walls D, Redmond EM, Cahill PA. Sonic Hedgehog Induces Notch Target Gene Expression in Vascular Smooth Muscle Cells via VEGF-A. Arterioscler Thromb Vasc Biol. American Heart Association; 2009 July 1;29(7):1112–1118.

26. Hulin-Curtis S, Williams H, Wadey KS, Sala-Newby GB, George SJ. Targeting Wnt/β-Catenin Activated Cells with Dominant-Negative N-cadherin to Reduce Neointima Formation. Mol Ther - Methods Clin Dev. 2017 June 16;5:191–199.

27. Riascos-Bernal DF, Chinnasamy P, Gross JN, Almonte V, Egaña-Gorroño L, Parikh D, Jayakumar S, Guo L, Sibinga NES. Inhibition of Smooth Muscle β-Catenin Hinders Neointima Formation After Vascular Injury. Arterioscler Thromb Vasc Biol. American Heart Association; 2017 May 1;37(5):879–888.

28. Zhu X, Wang Y, Soaita I, Lee HW, Bae H, Boutagy N, Bostwick A, Zhang RM, Bowman C, Xu Y, Trefely S, Chen Y, Qin L, Sessa W, Tellides G, Jang C, Snyder NW, Yu L, Arany Z, Simons M. Acetate controls endothelial-to-mesenchymal transition. Cell Metab. 2023 July 11;35(7):1163–1178.e10.

29. Chen S, He Q, Yang H, Huang H. Endothelial Birc3 promotes renal fibrosis through modulating Drp1-mediated mitochondrial fission via MAPK/PI3K/Akt pathway. Biochem Pharmacol. 2024 Nov 1;229:116477.

30. Zhang H, Tsui CK, Garcia G, Joe LK, Wu H, Maruichi A, Fan W, Pandovski S, Yoon PH, Webster BM, Durieux J, Frankino PA, Higuchi-Sanabria R, Dillin A. The extracellular matrix integrates mitochondrial homeostasis. Cell. Elsevier; 2024 Aug 8;187(16):4289–4304.e26.

31. Caruso P, Dunmore BJ, Schlosser K, Schoors S, Dos Santos C, Perez-Iratxeta C, Lavoie JR, Zhang H, Long L, Flockton AR, Frid MG, Upton PD, D’Alessandro A, Hadinnapola C, Kiskin FN, Taha M, Hurst LA, Ormiston ML, Hata A, Stenmark KR, Carmeliet P, Stewart DJ, Morrell NW. Identification of MicroRNA-124 as a Major Regulator of Enhanced Endothelial Cell Glycolysis in Pulmonary Arterial Hypertension via PTBP1 (Polypyrimidine Tract Binding Protein) and Pyruvate Kinase M2. Circulation. American Heart Association; 2017 Dec 19;136(25):2451–2467.

32. Uehara S, Chase CM, Kitchens WH, Rose HS, Colvin RB, Russell PS, Madsen JC. NK Cells Can Trigger Allograft Vasculopathy: The Role of Hybrid Resistance in Solid Organ Allografts1. J Immunol. 2005 Sept 1;175(5):3424–3430.

33. Franco-Acevedo A, Pathoulas CL, Murphy PA, Valenzuela NM. The Transplant Bellwether: Endothelial Cells in Antibody-Mediated Rejection. J Immunol. 2023 Nov 1;211(9):1276–1285.

34. Cross AR, Glotz D, Mooney N. The Role of the Endothelium during Antibody-Mediated Rejection: From Victim to Accomplice. Front Immunol [Internet]. 2018;Volume 9-2018. Available from: https://www.frontiersin.org/journals/immunology/articles/10.3389/fimmu.2018.00106

35. Watson CJ, Collier P, Tea I, Neary R, Watson JA, Robinson C, Phelan D, Ledwidge MT, McDonald KM, McCann A, Sharaf O, Baugh JA. Hypoxia-induced epigenetic modifications are associated with cardiac tissue fibrosis and the development of a myofibroblast-like phenotype. Hum Mol Genet. 2014 Apr 15;23(8):2176–2188.

36. Abe H, Takeda N, Isagawa T, Semba H, Nishimura S, Morioka MS, Nakagama Y, Sato T, Soma K, Koyama K, Wake M, Katoh M, Asagiri M, Neugent ML, Kim J whan, Stockmann C, Yonezawa T, Inuzuka R, Hirota Y, Maemura K, Yamashita T, Otsu K, Manabe I, Nagai R, Komuro I. Macrophage hypoxia signaling regulates cardiac fibrosis via Oncostatin M. Nat Commun. 2019 June 27;10(1):2824.

37. Gramley F, Lorenzen J, Pezzella F, Kettering K, Himmrich E, Plumhans C, Koellensperger E, Munzel T. Hypoxia and Myocardial Remodeling in Human Cardiac Allografts: A Time-course Study. J Heart Lung Transplant. Elsevier; 2009 Nov 1;28(11):1119–1126.

38. Zhao Y, Xiong W, Li C, Zhao R, Lu H, Song S, Zhou Y, Hu Y, Shi B, Ge J. Hypoxia-induced signaling in the cardiovascular system: pathogenesis and therapeutic targets. Signal Transduct Target Ther. 2023 Nov 20;8(1):431.

39. Daud A, Xu D, Revelo MP, Shah Z, Drakos SG, Dranow E, Stoddard G, Kfoury AG, Hammond MEH, Nativi-Nicolau J, Alharethi R, Miller DV, Gilbert EM, Wever-Pinzon O, McKellar SH, Afshar K, Khan F, Fang JC, Selzman CH, Stehlik J. Microvascular Loss and Diastolic Dysfunction in Severe Symptomatic Cardiac Allograft Vasculopathy. Circ Heart Fail. American Heart Association; 2018 Aug 1;11(8):e004759.

40. Jansen MAA, Otten HG, de Weger RA, Huibers MMH. Immunological and Fibrotic Mechanisms in Cardiac Allograft Vasculopathy. Transplantation [Internet]. 2015;99(12). Available from: https://journals.lww.com/transplantjournal/fulltext/2015/12000/immunological_and_fibrotic_mechanisms_in_cardiac.12.aspx

41. Stoeckius M, Hafemeister C, Stephenson W, Houck-Loomis B, Chattopadhyay PK, Swerdlow H, Satija R, Smibert P. Simultaneous epitope and transcriptome measurement in single cells. Nat Methods. United States; 2017 Sept;14(9):865–868. PMCID: PMC5669064

42. Tellides G, Tereb DA, Kirkiles-Smith NC, Kim RW, Wilson JH, Schechner JS, Lorber MI, Pober JS. Interferon-γ elicits arteriosclerosis in the absence of leukocytes. Nature. 2000 Jan 1;403(6766):207–211.

43. Nagano H, Mitchell RN, Taylor MK, Hasegawa S, Tilney NL, Libby P. Interferon-gamma deficiency prevents coronary arteriosclerosis but not myocardial rejection in transplanted mouse hearts. J Clin Invest. The American Society for Clinical Investigation; 1997 Aug 1;100(3):550–557.

44. Tellides G, Pober JS. Interferon-γ Axis in Graft Arteriosclerosis. Circ Res. American Heart Association; 2007 Mar 16;100(5):622–632.

45. Aguilar V, Le Master E, Paul A, Ahn SJ, Lazarko D, Febbraio M, Mehta D, Lee JC, Levitan I. Endothelial Stiffening Induced by CD36-Mediated Lipid Uptake Leads to Endothelial Barrier Disruption and Contributes to Atherosclerotic Lesions. Arterioscler Thromb Vasc Biol. American Heart Association; 2025 June 1;45(6):e201–e216.

46. Kim I doo, Ju H, Minkler J, Jiang R, Singh A, Sharma R, Febbraio M, Cho S. Endothelial cell CD36 mediates stroke-induced brain injury via BBB dysfunction and monocyte infiltration in normal and obese conditions. J Cereb Blood Flow Metab. SAGE Publications Ltd STM; 2023 June 1;43(6):843–855.

47. Chen Y, Yang M, Huang W, Chen W, Zhao Y, Schulte ML, Volberding P, Gerbec Z, Zimmermann MT, Zeighami A, Demos W, Zhang J, Knaack DA, Smith BC, Cui W, Malarkannan S, Sodhi K, Shapiro JI, Xie Z, Sahoo D, Silverstein RL. Mitochondrial Metabolic Reprogramming by CD36 Signaling Drives Macrophage Inflammatory Responses. Circ Res. American Heart Association; 2019 Dec 6;125(12):1087–1102.

48. Zhang Q, Li J, Liu X, Chen X, Zhu L, Zhang Z, Hu Y, Zhao T, Lou H, Xu H, Zhao W, Dong X, Sun Z, Sun X, Yang B, Zhang Y. Inhibiting CD36 palmitoylation improves cardiac function post-infarction by regulating lipid metabolic homeostasis and autophagy. Nat Commun. 2025 July 17;16(1):6602.

49. Lichscheidt ED, Jespersen NR, Nielsen BRR, Berg K, Seefeldt J, Nyengaard JR, Bøtker HE, Eiskjær H. Abnormal mitochondrial function and morphology in heart transplanted patients with cardiac allograft vasculopathy. J Heart Lung Transplant. 2022 June 1;41(6):732–741.

50. Monteiro JP, Rodor J, Caudrillier A, Scanlon JP, Spiroski AM, Dudnakova T, Pflüger-Müller B, Shmakova A, von Kriegsheim A, Deng L, Taylor RS, Wilson-Kanamori JR, Chen SH, Stewart K, Thomson A, Mitić T, McClure JD, Iynikkel J, Hadoke PWF, Denby L, Bradshaw AC, Caruso P, Morrell NW, Kovacic JC, Ulitsky I, Henderson NC, Caporali A, Leisegang MS, Brandes RP, Baker AH. MIR503HG Loss Promotes Endothelial-to-Mesenchymal Transition in Vascular Disease. Circ Res. American Heart Association; 2021 Apr 16;128(8):1173–1190.

51. Weigand JE, Boeckel JN, Gellert P, Dimmeler S. Hypoxia-Induced Alternative Splicing in Endothelial Cells. PLOS ONE. Public Library of Science; 2012 Aug 2;7(8):e42697.

52. Raghunandan S, Ramachandran S, Ke E, Miao Y, Lal R, Chen ZB, Subramaniam S. Heme Oxygenase-1 at the Nexus of Endothelial Cell Fate Decision Under Oxidative Stress. Front Cell Dev Biol [Internet]. 2021;Volume 9-2021. Available from: https://www.frontiersin.org/journals/cell-and-developmental-biology/articles/10.3389/fcell.2021.702974

53. Tran DT, Esckilsen S, Mulligan J, Mehrotra S, Atkinson C, Nadig SN. Impact of Mitochondrial Permeability on Endothelial Cell Immunogenicity in Transplantation. Transplantation [Internet]. 2018;102(6). Available from: https://journals.lww.com/transplantjournal/fulltext/2018/06000/impact_of_mitochondrial_permeability_on.16.aspx

54. Chen M, Zhang J, Manley JL. Turning on a Fuel Switch of Cancer: hnRNP Proteins Regulate Alternative Splicing of Pyruvate Kinase mRNA. Cancer Res. 2010 Nov 14;70(22):8977–8980.

55. Xu Y, Ma X, Ni W, Zheng L, Lin Z, Lai Y, Yang N, Dai Z, Yao T, Chen Z, Shen L, Wang H, Wang L, Wu Y, Gao W. PKM2-Driven Lactate Overproduction Triggers Endothelial-To-Mesenchymal Transition in Ischemic Flap via Mediating TWIST1 Lactylation. Adv Sci. John Wiley & Sons, Ltd; 2024 Dec 1;11(47):2406184.

56. Marques E, Kramer R, Ryan DG. Multifaceted mitochondria in innate immunity. Npj Metab Health Dis. 2024 May 27;2(1):6.

57. Huang LS, Hong Z, Wu W, Xiong S, Zhong M, Gao X, Rehman J, Malik AB. mtDNA Activates cGAS Signaling and Suppresses the YAP-Mediated Endothelial Cell Proliferation Program to Promote Inflammatory Injury. Immunity. Elsevier; 2020 Mar 17;52(3):475–486.e5.

58. Hanada Y, Maeda R, Ishihara T, Nakahashi M, Matsushima Y, Ogasawara E, Oka T, Ishihara N. Alternative splicing of Mff regulates AMPK-mediated phosphorylation, mitochondrial fission and antiviral response. Pharmacol Res. 2024 Nov 1;209:107414.

59. Rockfield SM, Turnis ME, Rodriguez-Enriquez R, Bathina M, Ng SK, Kurtz N, Becerra Mora N, Pelletier S, Robinson CG, Vogel P, Opferman JT. Genetic ablation of *Immt* induces a lethal disruption of the MICOS complex. Life Sci Alliance. 2024 June 1;7(6):e202302329.

60. Gomes-Silva B, Furtado M, Ribeiro M, Martins S, Carvalho MT, Maatz H, Radke M, Gotthardt M, Savisaar R, Carmo-Fonseca M. Alternative splicing dynamics during human cardiac development *in vivo* and *in vitro*. bioRxiv. 2025 Jan 1;2025.03.21.642423.

61. Zhu C, Wu J, Sun H, Briganti F, Meder B, Wei W, Steinmetz LM. Single-molecule, full-length transcript isoform sequencing reveals disease-associated RNA isoforms in cardiomyocytes. Nat Commun. 2021 July 9;12(1):4203.

62. Hong SG, Shin J, Choi SY, Powers JC, Meister BM, Sayoc J, Son JS, Tierney R, Recchia FA, Brown MD, Yang X, Park JY. Flow pattern–dependent mitochondrial dynamics regulates the metabolic profile and inflammatory state of endothelial cells. JCI Insight [Internet]. The American Society for Clinical Investigation; 2022 Sept 22;7(18). Available from: 10.1172/jci.insight.159286

63. Tran DT, Tu Z, Alawieh A, Mulligan J, Esckilsen S, Quinn K, Sundararaj K, Wallace C, Finnegan R, Allen P, Mehrotra S, Atkinson C, Nadig SN. Modulating donor mitochondrial fusion/fission delivers immunoprotective effects in cardiac transplantation. Am J Transplant Off J Am Soc Transplant Am Soc Transpl Surg. 2022 Feb;22(2):386–401. PMCID: PMC8813895

64. Colucci F, Di Santo JP. The receptor tyrosine kinase c-kit provides a critical signal for survival, expansion, and maturation of mouse natural killer cells. Blood. 2000 Feb 1;95(3):984–991.

65. Schierloh P, Yokobori N, Geffner L, Balboa L, Romero MM, Musella RM, Alemán M, Castagnino J, Basile J, de la Barrera SS, Abbate E, Sasiain MC. NK cells from tuberculous pleurisy express high ICAM-1 levels and exert stimulatory effect on local T cells. Eur J Immunol. John Wiley & Sons, Ltd; 2009 Sept 1;39(9):2450–2458.

66. Aziz T, Hasleton P, Hann AW, Yonan N, Deiraniya A, Hutchinson IV. Transforming growth factor beta in relation to cardiac allograft vasculopathy after heart transplantation. J Thorac Cardiovasc Surg. 2000 Apr;119(4 Pt 1):700–708. PMID: 10733758

67. Labarrere CA, Woods JR, Hardin JW, Campana GL, Ortiz MA, Jaeger BR, Reichart B, Bonnin JM, Currin A, Cosgrove S, Pitts DE, Kirlin PC, O’Donnell JA, Hormuth DA, Wozniak TC. Early prediction of cardiac allograft vasculopathy and heart transplant failure. Am J Transplant Off J Am Soc Transplant Am Soc Transpl Surg. 2011 Mar;11(3):528–535. PMID: 21219580

68. Labelle M, Begum S, Hynes RO. Direct signaling between platelets and cancer cells induces an epithelial-mesenchymal-like transition and promotes metastasis. Cancer Cell. 2011 Nov 15;20(5):576–590. PMCID: PMC3487108

69. Sun H, Schlamp F, Muller M, Xia Y, Liberow S, Smilowitz NR, Hochman JS, Reynolds HR, Beckman JA, Barrett TJ, Berger JS. Platelets induce endothelial cell mitochondrial dysfunction in myocardial infarction. Sci Adv. American Association for the Advancement of Science; 11(46):eadx0268.

70. Murphy PA, Butty VL, Boutz PL, Begum S, Kimble AL, Sharp PA, Burge CB, Hynes RO. Alternative RNA splicing in the endothelium mediated in part by Rbfox2 regulates the arterial response to low flow. Elife. 2018 Jan 2;7. PMCID: PMC5771670

71. Chen Z, He C, Gao Z, Li Y, He Q, Wang Y, Cai C. Polypyrimidine tract binding protein 1 exacerbates cardiac fibrosis by regulating fatty acid-binding protein 5. ESC Heart Fail. John Wiley & Sons, Ltd; 2023 June 1;10(3):1677–1688.

72. Tseng PL, Sun W, Salem A, Alaklobie M, Macfarlane SC, Gad AKB, Collins MO, Erdmann KS. Mechanical control of the alternative splicing factor PTBP1 regulates extracellular matrix stiffness induced proliferation and cell spreading. iScience [Internet]. Elsevier; 2025 Apr 18 [cited 2026 Feb 12];28(4). Available from: 10.1016/j.isci.2025.112273

73. Tran DT, Esckilsen S, Mulligan J, Mehrotra S, Atkinson C, Nadig SN. Impact of Mitochondrial Permeability on Endothelial Cell Immunogenicity in Transplantation. Transplantation [Internet]. 2018;102(6). Available from: https://journals.lww.com/transplantjournal/fulltext/2018/06000/impact_of_mitochondrial_permeability_on.16.aspx

74. Mohanta SK, Heron C, Klaus-Bergmann A, Horstmann H, Brakenhielm E, Giannarelli C, Habenicht AJR, Gerhardt H, Weber C. Metabolic and Immune Crosstalk in Cardiovascular Disease. Circ Res. American Heart Association; 2025 May 23;136(11):1433–1453.

75. Eelen G, de Zeeuw P, Simons M, Carmeliet P. Endothelial Cell Metabolism in Normal and Diseased Vasculature. Circ Res. American Heart Association; 2015 Mar 27;116(7):1231–1244.

76. Li X, Fang P, Yang WY, Chan K, Lavallee M, Xu K, Gao T, Wang H, Yang X. Mitochondrial ROS, uncoupled from ATP synthesis, determine endothelial activation for both physiological recruitment of patrolling cells and pathological recruitment of inflammatory cells. Can J Physiol Pharmacol. NRC Research Press; 2017 Mar 1;95(3):247–252.

77. Zhu X, Wang Y, Soaita I, Lee HW, Bae H, Boutagy N, Bostwick A, Zhang RM, Bowman C, Xu Y, Trefely S, Chen Y, Qin L, Sessa W, Tellides G, Jang C, Snyder NW, Yu L, Arany Z, Simons M. Acetate controls endothelial-to-mesenchymal transition. Cell Metab. 2023 July 11;35(7):1163–1178.e10.

78. Sanders E, Kaur K, Wagner N, Emig R, Aronovitz M, Bayer AL, Theall B, Tai A, Martino N, Good ME, Adam AP, Blanton R, Alcaide P. Endothelial Cell Stimulator of Interferon Genes Regulates IL-6 Production and Is Required for Pathologic Cardiac Hypertrophy and Contractile Dysfunction in Experimental Heart Failure. Am J Pathol. 2025 Oct;195(10):1788–1807. PMCID: PMC12597551

79. Clerkin KJ, Topkara VK, Farr MA, Jain R, Colombo PC, Restaino S, Sayer G, Castillo M, Lam EY, Chernovolenko M, Yuzefpolskaya M, DeFilippis E, Latif F, Zorn E, Takeda K, Johnson LL, Uriel N, Einstein AJ. Noninvasive Physiologic Assessment of Cardiac Allograft Vasculopathy Is Prognostic for Post-Transplant Events. J Am Coll Cardiol. 2022 Oct 25;80(17):1617–1628. PMCID: PMC9758655

80. Hensel JA, Nicholas SAE, Kimble AL, Nagpal AS, Omar OMF, Tyburski JD, Jellison ER, Ménoret A, Ozawa M, Rodriguez-Oquendo A, Vella AT, Murphy PA. Splice factor polypyrimidine tract-binding protein 1 (Ptbp1) primes endothelial inflammation in atherogenic disturbed flow conditions. Proc Natl Acad Sci. Proceedings of the National Academy of Sciences; 2022 July 26;119(30):e2122227119.

81. Corry RJ, Winn HJ, Russell PS. Primarily vascularized allografts of hearts in mice. The role of H-2D, H-2K, and non-H-2 antigens in rejection. Transplantation. 1973 Oct;16(4):343–350. PMID: 4583148

82. Li YI, Knowles DA, Humphrey J, Barbeira AN, Dickinson SP, Im HK, Pritchard JK. Annotation-free quantification of RNA splicing using LeafCutter. Nat Genet. 2018 Jan;50(1):151–158. PMCID: PMC5742080

